# Inference on paths of yeast transcription pre-initiation complex assembly by model-based analysis of altered occupancy profiles

**DOI:** 10.64898/2026.09.30.755594

**Authors:** Elizaveta A. Novikova, Arach Goldar, Pierre Nicolas, Julie Soutourina

**Affiliations:** Université Paris-Saclay, CEA, CNRS, Institute for Integrative Biology of the Cell (I2BC), 91198, Gif-sur-Yvette, France, Equipe Labellisée LIGUE Contre le Cancer; Université Paris-Saclay, INRAE, MaIAGE, Jouy-en-Josas, France

## Abstract

Dysfunctions in transcription cause severe human diseases. In eukaryotes, transcription initiation by RNA polymerase (Pol) II requires the formation of the pre-initiation complex (PIC) composed of general transcription factors (GTFs), Pol II, and an essential multiprotein coactivator, Mediator. *In vitro* experiments with GTFs and Pol II have depicted a linear sequence of PIC assembly. However, PIC assembly paths *in vivo* at the genomic scale and Mediator’s role in these paths remain poorly understood. One strategy characterizes ChIP-seq occupancy alterations in Mediator mutants, but analysis of these data still lacks a formalized interpretation framework. In this study, we address this need by developing a quantitative framework that scores candidate assembly paths according to their ability to explain these alterations. This framework combines a model of PIC assembly based on a system of differential equations with parameter optimization based on a penalized criterion. We applied this new framework to datasets from two yeast Mediator mutants, in the Med17 and Med10 subunits. Our results show that the best-supported PIC assembly paths differ from those inferred *in vitro*. In particular, we show that it is important to distinguish the core TFIIH module from the TFIIK kinase module, and that similar paths starting with Mediator or TBP receive different levels of support. Furthermore, although multiple paths fit the data equally well, they differ markedly for specific gene groups. Taken together, the results show that the path can depend on the gene, and are also compatible with more than one path being operational *in vivo* for a single gene. This work improves our understanding of PIC assembly paths and transcription initiation, and opens perspectives for future modeling and experimental studies.

## 1 Introduction

Transcription, the first step of gene expression, is fundamental to all living organisms, and its dysfunction underlies serious human diseases, such as neurodevelopmental pathologies and cancers. In eukaryotes, RNA polymerases (Pol) II transcribe all protein-coding genes and a large variety of non-coding RNA genes. Pol II, as the other eukaryotic RNA polymerases, is unable to recognize gene promoters without the assistance of the general transcription factors (GTFs): TFIIA, TFIIB, TFIID, TFIIE, TFIIF and TFIIH, named from the protein fractionation of nuclear extracts that allowed purification of these factors (reviewed in [1, 2]). The GTFs and Pol II assemble *in vitro* on promoters to form the pre-initiation complex (PIC) and initiate transcription at a basal level. Textbook models of transcription initiation based on *in vitro* biochemical experiments depict a linear sequence of assembly of the GTFs and Pol II in PIC. This assembly starts with the recruitment of TATA-binding protein (TBP)-containing TFIID on promoter DNA, followed by TFIIA, TFIIB, Pol II together with TFIIF, and then TFIIH along with TFIIE, completing the PIC [3, 4]. Of note, these studies did not distinguish core TFIIH and its kinase module, TFIIK.

Mediator was not taken into account in these initial experiments despite the importance of this complex for PIC assembly *in vitro* and *in vivo* (reviewed in [5, 6]). Indeed, Mediator is an essential multiprotein coactivator that serves as a link between the transcription activators and the basal transcription machinery for transcription activation and promotes the PIC assembly. Mediator is involved in physical and functional interactions with many PIC components including Pol II, TFIID, TFIIH and TFIIB [7]. Further biochemical experiments have indicated that *in vitro* the first round of transcription is slower than the subsequent reinitiation rounds. The fast reinitiation step is stimulated by a scaffold complex, consisting of Mediator and several GTFs such as TFIIA, TFIID, TFIIH and TFIIE, which remain associated with promoters after Pol II has initiated transcription [8]; reinitiation remains to be demonstrated *in vivo*. Recent development in cryoEM allowed to obtain a global view of PIC and resulted in a structure of entire human PIC including TFIID and Mediator [9] or yeast PIC engaged with the +1 nucleosome [10].

In order to understand PIC formation and transcription initiation in the cellular context, including the presence of Mediator and promoter diversity, it is necessary to investigate PIC assembly paths *in vivo*. The order of PIC component recruitment has first been investigated for inducible promoters in yeast *Saccharomyces cerevisiae*, demonstrating that, *in vivo*, Mediator was recruited before the GTFs and Pol II [11–13]. Further *in vivo* studies, with yeast Mediator mutants, led to propose a model of PIC assembly in which Mediator independently orchestrates several steps of this process [7, 14, 15]. In particular, ChIP-seq experiments allowed to look for changes in the recruitment of Mediator, GTFs and Pol II at genomic scale in these Mediator mutants, and the diversity of alteration profiles in PIC component occupancy suggested that the assembly step that was affected and/or the assembly paths could differ depending on the promoter. Diversity and complexity in PIC assembly was supported by several studies and a partial PIC was proposed as a regulated intermediate [5, 16, 17]. Moreover, recent studies of TFIIB or TBP further challenged a classical model of a stable sequential binding of protein factors to promoters for PIC assembly and suggested that the interactions of PIC components with promoter DNA are more transient and dynamic, highlighting the complexity of the assembly process [18–21]. Using single-molecule tracking in live yeast cells, a recent study emphasized a dynamic PIC assembly and a key role of TFIID/TBP, Mediator and Pol II in coordinating this process [22]. With regards to Mediator, interactions of this complex with PIC components on core promoters are thought to be also transient which potentially contributes to accommodate multiple contacts while avoiding clashes [6, 23, 24]. Taken together, *in vivo* studies suggest more complex relationships between PIC components than the linear assembly model initially proposed. It should be noted that a recent single-molecule *in vitro* study with yeast nuclear extracts also argues against a simple sequential PIC assembly, proposing instead more complex activator-dependent assembly paths [25].

Mathematical models are important for connecting quantitative data and qualitative biological hypotheses. Assembly of multiprotein complexes involved in transcription or DNA repair was studied *in silico* by combining analyses of live-cell imaging kinetics and mathematical modeling via differential equations (reviewed in [26]). A modeling approach for gene transcription based on the *in situ* hybridization data for visualization of expression level of developmental control genes was proposed to analyze the impact of cell-to-cell variations, RNA polymerase loading, promoter-proximal pausing and elongation on mRNA expression [27]. In this study, PIC assembly was only included as one of the regulation step without any specific modeling. A simulation program to evaluate allowable disassembly paths in the context of a fixed PIC assembly path was developed based on ChIP-chip data for PIC components including Pol II and GTFs in a wild-type yeast strain [28]. Although they provided very valuable insights, analyses of PIC component occupancy profiles in mutants by ChIP-seq and addressing the probability of different PIC assembly paths lacked a formal modeling framework.

In this study, we developed a modeling framework for the quantitative analysis of mutant vs. wildtype ChIP-seq data in the context of PIC assembly studies. This approach, which combines modeling PIC assembly with a system of differential equations and parameter optimization based on a penalized criterion, allows to score candidate PIC assembly paths according to their ability to explain alteration profiles in PIC component occupancy. We re-examine with this new methodological framework datasets available for two Mediator mutants: *med17-504* in Med17 subunit and *med10-196* in Med10 subunit. Based on protein-protein interaction data, these two mutants were previously reported, respectively, as affected in Mediator-Pol II interaction [15] and Mediator-TFIIB interaction [7]. The joint analysis of these two ChIP-seq datasets allows to identify the best-supported PIC assembly paths and to simultaneously obtain information on which PIC assembly steps are altered in each Mediator mutant.

## 2 Data, model and inference

### 2.1 Experimental data

ChIP-seq allows genome-wide profiling of the population averaged instantaneous frequency of the presence of a protein along the genome. Direct quantitative comparison between proteins is complicated by variable efficiency of cross-linking and pull-down, which induces differences in the scale of the occupancy profiles. However, quantitative comparison of occupancy profiles for a same protein between two genetic backgrounds is not affected by these protein-dependent variations and proved useful to establish functional links between PIC components. Here, we used ChIP-seq tag density profiles collected for this purpose on two mutants, *med17-504* [15] and *med10-196* [7]. These profiles consisted of normalized number of reads, extended to average DNA-fragment length, overlapping each position of the genome.

In keeping with [7, 15], tag density in promoters was averaged at the promoter level and the impact of each mutation on occupancy of 5 to 6 PIC components was quantified by computing ratios between mutant and wild-type values. For TBP, Mediator represented by its Med15 subunit, TFIIB (not available for *med17-504*), TFIIK, and core TFIIH, tag densities were averaged across whole intergenic regions between ORFs of tandem genes. For Pol II, which is recruited at the TSS during PIC assembly but which is also found over the whole transcript during elongation, tag density was averaged on a more restricted region of length 100 bp directly upstream the TSS. Divergently transcribed genes and class III genes were excluded from the analysis. Regions with a null tag density for a PIC component in mutant or wild-type backgrounds in any of two studies were also excluded. The per-gene mutant vs. wild-type ratios are provided in Supplementary Table S1.

These ratios were aggregated (by computing the median) for the entire gene set for 16 subsets intended to account for the diversity of promoters (Fig. 1). A total of 7 subsets were defined based on prior knowledge of biologically-relevant characteristics or transcription level (Fig. 1A): (i-ii) presence or absence of TATA-box, (iii) genes encoding ribosomal proteins, (iv) genes belonging to ribosome biogenesis (Ribi) regulon [29], or (v-viii) according to transcription level as measured by Pol II occupancy in wild-type averaged across the entire ORF averaged between both studies (3 groups were distinguished, corresponding respectively to Pol II tag density in the lower quartile, in the interquartile range, in the upper quartile). Furthermore, we considered also the 9 subsets defined in [7] (Fig. 1B). These correspond to strong, intermediate, and weak effect of the *med10-196* mutation on TFIIB, Med17, and Pol II occupancy, with each time the intermediate subset being twice larger than the two others (genes with low transcription levels were excluded from the definition of these subsets). The list of genes constitutive of each set is found in Supplementary Table S2 and the numerical values of the mutant vs. wild-type ratios are given in Supplementary Table S3.

**Figure 1:**
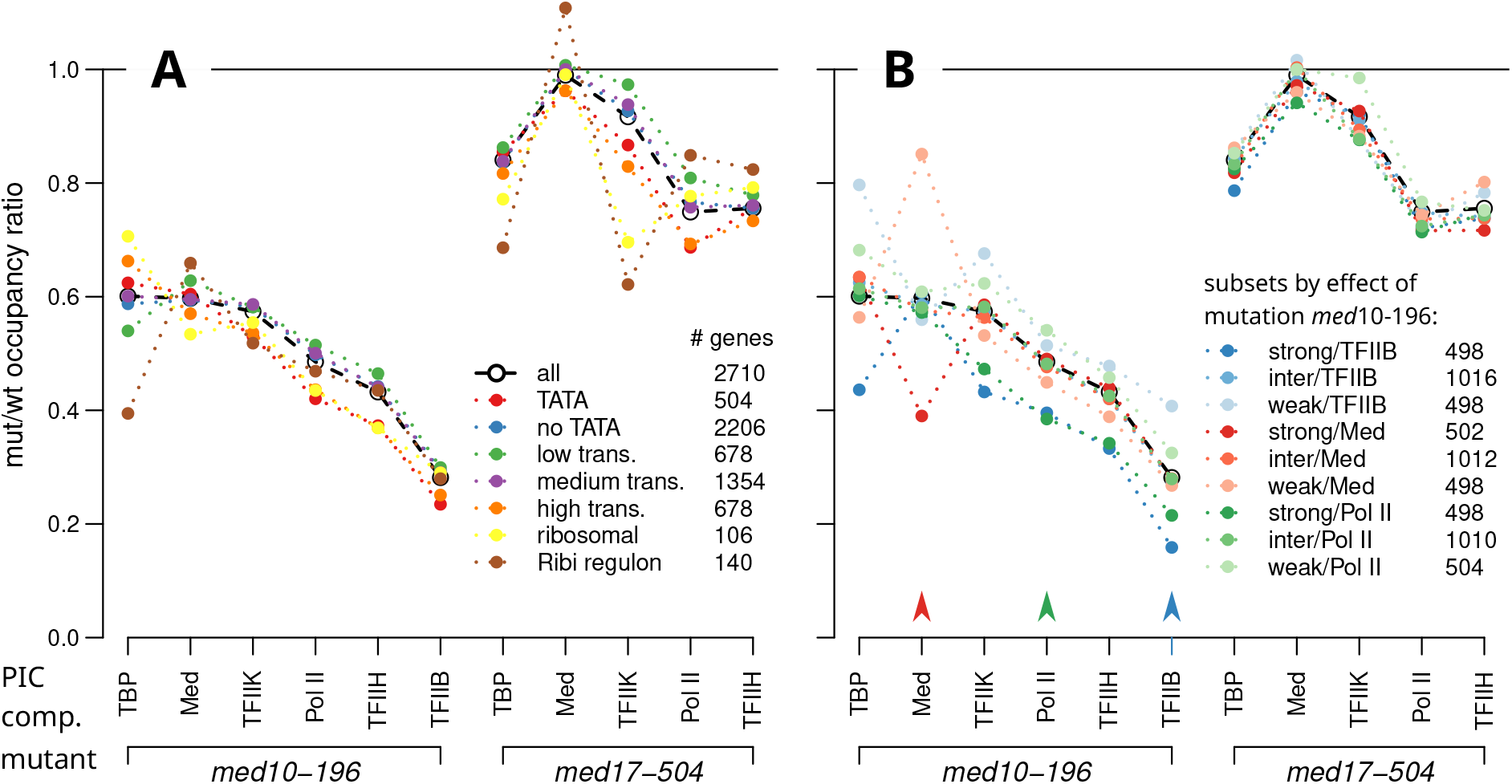
Mutant vs. wild-type ratios of occupancy by PIC components in two mutants for 17 sets of genes. The median ratio measured by ChIP-seq is shown for the entire gene set (dataset “all”) and different subsets. The PIC components are ordered on the x-axis to display the ratios for the entire set in the *med10-196* mutant by decreasing values. **A** The 7 subsets of genes distinguished by prior knowledge of biologically-relevant characteristics or transcription level. **B** The 9 subsets of genes distinguished by alteration (strongest quartile, intermediate, weakest quartile) in the *med10-196* mutant for three PIC components; arrows highlight the PIC components taken into account to define these groups (Med, Pol II and TFIIB).

### 2.2 A mechanistic model of PIC assembly in wild-type and mutant

The focus of the inference is to identify the sequence of recruitment, or “assembly path”, of the 6 major PIC components studied by ChIP-seq. To be parsimonious, we base this inference on a minimal model of PIC assembly that describes only the transitions between 7 PIC states corresponding to the presence, at a promoter, of 0 to 6 of these PIC components recruited along a fixed but unknown linear assembly path. Transitions are modeled using a set of kinetic rates, denoted *k*, which are altered in the mutants by some coefficients, denoted *a*. Below, we present the details of this dynamical model and explain how its predictions are compared to the observed data.

In the model, the population-averaged fraction of cells containing exactly *i* out of the 6 PIC components at a given promoter, denoted *x*_*i*_, results from the steady-state of a dynamical process involving 6 recruitment steps. Each of these individual steps is reversible but the completion of the PIC (state *i* = 6), permits the initiation of transcription by Pol II which is not reversible and leads to total or partial disassembly; the PIC returns to a state, without Pol II, in the linear assembly path (Fig. 2A). This writes as a system of first order ordinary differential equations,

**Figure 2:**
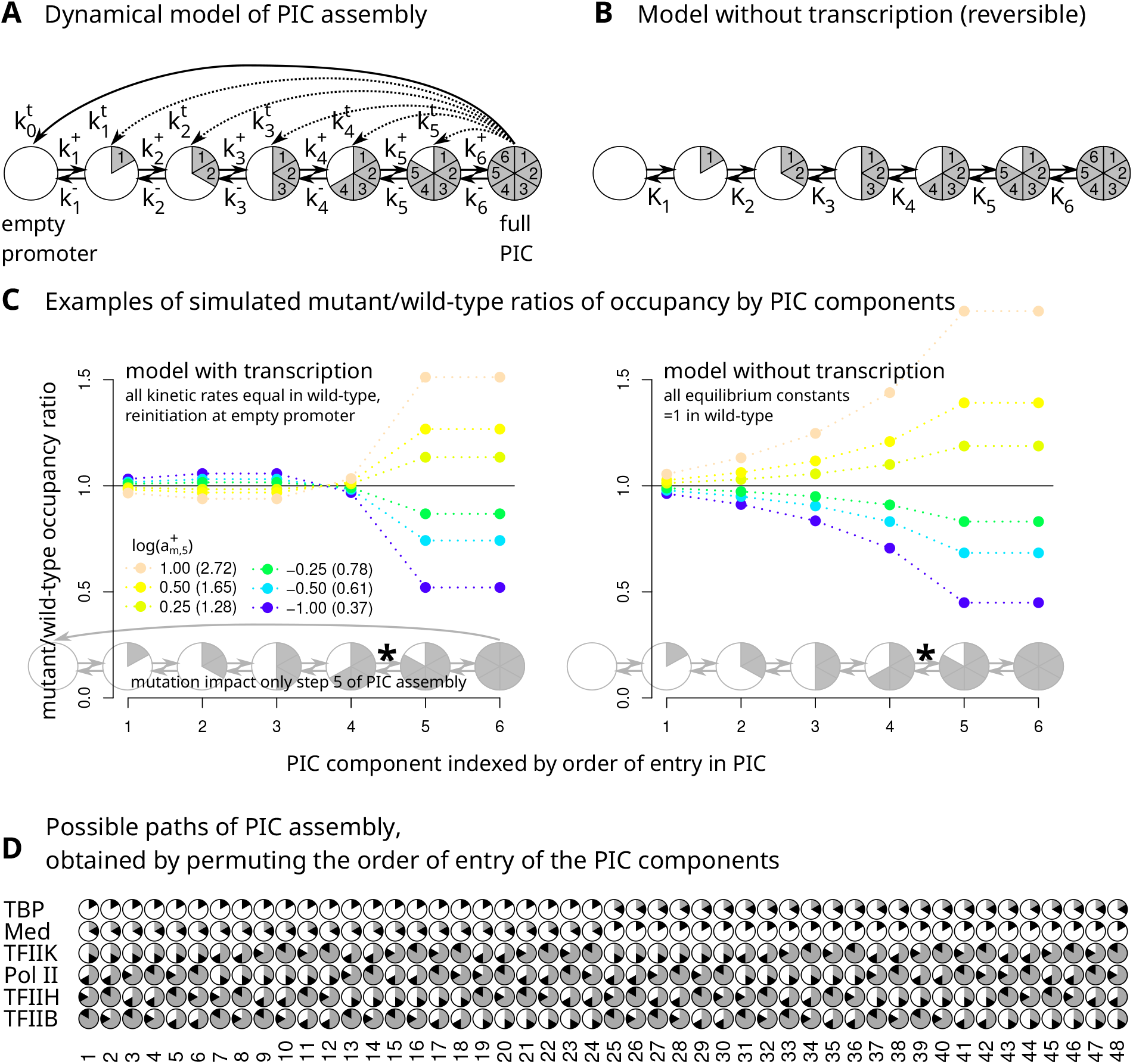
Modeling approach. **A** Dynamical model of PIC assembly. The degree of completeness of the PIC assembly is represented as a pie chart with 6 slices, each one corresponding to one of the PIC components considered in the study. Notations distinguish three sets of kinetic rates: assembly rates 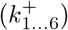, disassembly rates 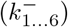, reinitiation rates when transcription starts 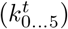. **B** When transcription is not taken into account, model is reversible and thus depends only on equilibrium constants (*K*_1…6_). **C** Examples of simulated PIC component mutant/wild-type occupancy ratios when only one step of the PIC assembly is altered by the mutation and all kinetic rates are set equal in wild-type. The color code reflects the alteration coefficient 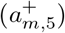 on the assembly rate of the 5^th^ step of PIC assembly 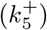, values are in log-scale (values in natural scale between parentheses). Left panel: model with transcription and reinitiation at empty promoter. Right panel: model without transcription. **D** The 48 possible PIC assembly paths analyzed in this study. For each path, the recruitment step of a given PIC component is represented by a circular diagram with 6 sectors filled clockwise, the sector corresponding to that component’s step highlighted in black.

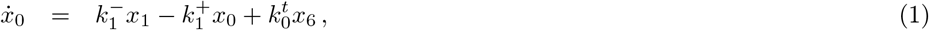

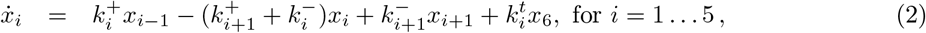

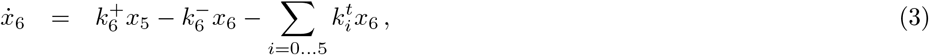

where 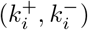 is the pair assembly/disassembly rates associated with the PIC assembly step *i*, and 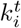 is the rate of transcription leading to reinitiation of the PIC assembly process at PIC state *i*. Solving, under the constraint ∑_*i*=0…6_ *x*_*i*_ = 1, the system of linear equations which equates to zero all these time derivatives gives the steady-state values. In a mutant, *m*, any of the rates governing the PIC assembly/disassembly can be altered and we denote by 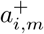 and 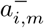 the alteration coefficients of assembly and disassembly rates of step *i, i*.*e* 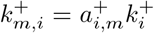 and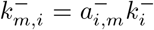. Given these rates and using the same approach as for *x*_*i*_, we obtain the fraction of PIC at state *i* in mutant *m*, denoted *x*_*i,m*_.

The experimental data take the form of mutant vs. wild-type occupancy ratios. Having numbered the PIC components according to their order of recruitment along the assembly path, the predicted mutant *vs*. wild-type ratio for the PIC component *i* writes

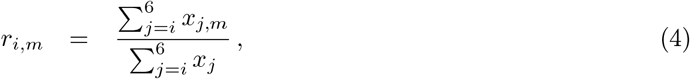

where the sums over *j* = *i* … 6 aggregate the predictions for the states containing the PIC component *i* in the mutant and in the wild-type.

Transcription involves the departure of Pol II from the promoter. After transcription initiation, PIC assembly thus resumes at a state before Pol II recruitment. Under the assumption of a simple linear path, if Pol II is recruited at step *l*the *l*^th^ component according to the the PIC assembly path, this implies that 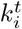 is allowed to be non-null only for *i < l*. To limit the number of free parameters, we only consider the possibility of reinitiation at the first or the last possible state, *i*.*e*. 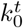 and 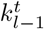 are allowed to be non-null. Accordingly, we distinguish four models that differ with respect to the consequences of transcription on the PIC assembly/disassembly process:

- negligible effect (model T_*∅*_, Fig. 2B);
- reinitiation at empty promoter (model T_1_);
- reinitiation at last PIC assembly state before recruitment of Pol II (model T_2_);
- reinitiation at both states (model T_1+2_).

In models T_1_, T_2_, and T_1+2_, when the kinetic rate of transcription initiation is comparable to the kinetic rates of the assembly/disassembly process, transcription substantially contributes to the shaping the predicted mutant vs. wild-type profiles (Fig. 2C).

Of note, in these models of the dynamical process of PIC assembly, distinct set parameter values predict the same mutant vs. wild-type occupancy profiles. For instance, if transcription has a negligible effect (model T_*∅*_, Fig. 2B), all PIC assembly steps are reversible, and thus only the 6 equilibrium constants 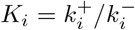 for *i* = 1 … 6 impact the global equilibrium occupancy profile of the wild-type. Similarly, in T_*∅*_, the 6 ratios 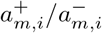, for *i* = 1 … 6, determine the occupancy profile of mutant *m* given the profile of the wild-type.

### 2.3 Scoring PIC assembly paths by penalized optimization

The aim here is to score the PIC assembly paths according to their ability to explain the experimental ChIP-seq data. This scoring involves numerical optimization of the parameters of the mechanistic model, (*k, a*), with respect to the distance between the predicted mutant vs. wild-type occupancy ratios, *r*(*k, a*), and their observed counterparts, denoted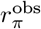. The symbol *π* denotes here the permutation which orders the occupancy ratios according to the sequence of recruitment of the components in the assembly process. Our search space consisted of the 48 permutations corresponding to all possible paths that start with either TBP or Med (Fig. 2D).

Not only different sets of parameter values can explain the same mutant vs. wild-type occupancy ratios for a given path, but also many paths can explain equally well the experimental data if no preference is given to simpler models. In this context, according to a parsimony principle (Occam’s razor), parameter sets and PIC assembly paths involving fewer alterations of assembly steps in the mutants should be preferred.

To implement this parsimony principle, we minimize, given the permutation *π*, a penalized objective function consisting of a sum of three terms,

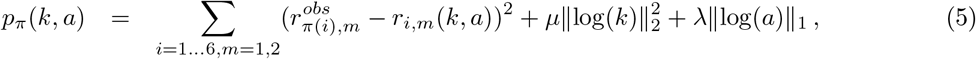

where *∥*log(*k*) *∥*_2_ and *∥*log(*a*)*∥*_1_ are shorthand notations for the 2-norm of the vector of natural logarithms of the kinetic rates and for the 1-norm (sum of absolute values) of the vector of natural logarithms of the coefficients of alteration in the two mutants, respectively. Each term has its specific purpose: the first is a sum of squared errors (SSE) that assesses the fit to the data; the second penalizes the dispersion of the kinetic rates and enforces the uniqueness of the best set of kinetic rate values; the third penalizes the number and amplitude of the alterations of the kinetic rates in the two mutants. In regularization theory, a 2-norm penalty is known as Tikhonov regularization [30] and the 1-norm penalty is known to induce sparsity [31]. Such penalty terms are used respectively in Ridge and Lasso regressions in the context of high-dimensional data analysis [32]. Here, they apply to distinct sets of parameters and their strengths are tuned independently by two coefficients *µ >* 0 and *λ >* 0. In practice, numerical optimization was carried out with Wolfram Mathematica (version 11.0.1) using the built-in function NMinimize (with default settings) and a symbolic expression for *r*(*k, a*).

The score of a path is the result of the above minimization for the corresponding permutation *π*,

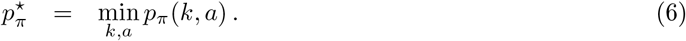

Paths can then be compared in terms of their relative score, *i*.*e*.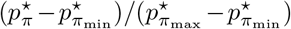, which yields a value between 0, for the best-scoring path (*π* = *π*_min_), and 1, for the worst-scoring path (*π* = *π*_max_).

## 3 Results and Discussion

### 3.1 Sensitivity and robustness with respect to penalty strengths

The penalized optimization scheme proposed in this work is based on a trade-off between achieving the fit to the observed mutant vs. wild-type profile (measured by the SSE) and minimizing two penalty terms. Penalty strengths increase with the values of two coefficients, *µ* and *λ*, which apply to the dispersion of the kinetic rates and to the amplitude of their alteration coefficients in the mutants, respectively. To explore how modifying the penalty strengths changes the results, we scored the 48 possible PIC assembly paths on the mutant vs. wild-type profile for the entire gene set, for each combination of *µ* and *λ* over a grid covering values from 10^*−*3^ to 1 (Fig. 3A). The general trends describing the respective impacts of *µ* and *λ* are similar from the model T_0_ (Fig. 3) to T_1+2_ (Supplementary Fig. S1).

**Figure 3:**
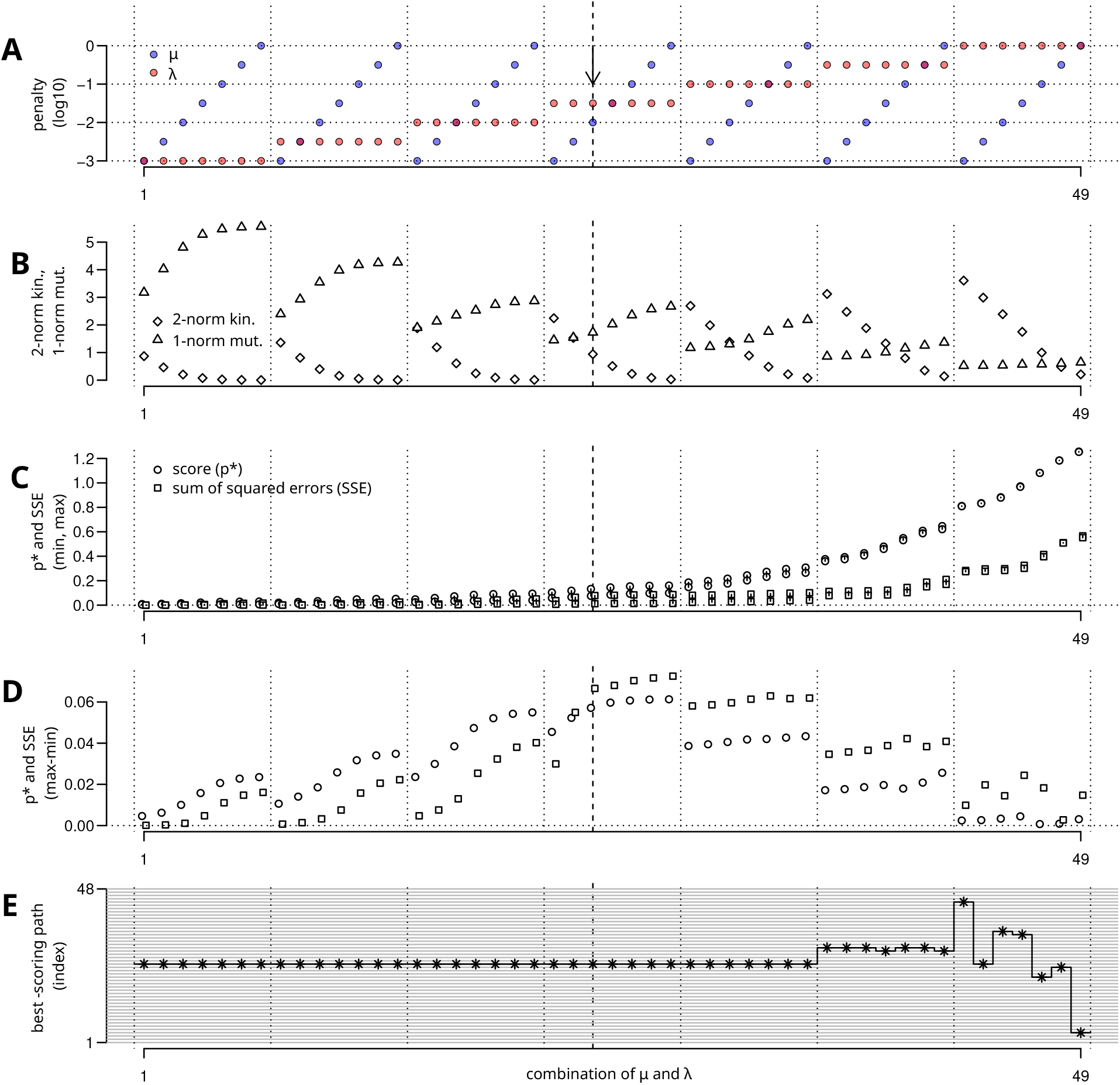
Varying penalty strengths on real data. Results obtained for 49 combinations of *µ* and *λ* values (index of combination on x-axis) with transcription model T_0_ on the entire gene set. **A** Values of penalty coefficients *µ* (blue points) and *λ* (red points). **B** Values of *∥*log(*k*) *∥*_2_ (diamonds) and *∥*log(*a*) *∥* _1_ (triangles), respectively the 2-norm involved in the penalization of the dispersion of the kinetic rates and the 1-norm involved in the penalization of the amplitude of the alteration coefficients in the mutants (Equation 5). These values are for the best-scoring path. **C** Minimum and maximum values for the score *p*^*⋆*^ (circles) and for the sum of squared errors of prediction (squares), across the 48 possible PIC assembly paths. **D** Difference between maximum and minimum values represented in **C. E** Index of the best-scoring path, *i*.*e*. the PIC assembly path minimizing the penalized optimization criterion *p*^*⋆*^. The vertical arrow in **A** and the dashed lines in **A-E** correspond to the position of the combination *λ* = 10^*−*1.5^ and *µ* = 10^*−*2^ that we selected for the inference.

Examining the parameters (*k, a*) associated with the best-scoring path confirms that increasing *µ* and *λ* drastically decreases the dispersion of the kinetic rates and the amplitude of the alteration coefficients in the mutants, respectively (Fig. 3B, see also Supplementary Fig. S2). We also note that, at fixed *λ*, increasing *µ* tends to increase the alteration coefficients in the mutants, probably in compensation to the drastic decrease of the dispersion of the kinetic rates.

Examining the fit to the data (in terms of SSE) across the whole grid reveals the preponderant role of *λ* (Fig. 3C). With a small value of *λ*, near 10^*−*3^, the optimized parameter sets perfectly explain the data, whatever the PIC assembly path, indicating that too much weight is given to the fit of the data. At the other end, a high value of *λ*, near 1, yields parameter sets that poorly explain the data, whatever the path, indicating that too much weight is given to minimizing the effect of the mutations. Between these two extremes are values of *λ*, near 10^*−*1.5^, for which the optimized parameter set depends on the PIC assembly path, both in terms of SSE and score *p*^*⋆*^(Fig. 3D). Like *λ*, decreasing *µ* decreases the SSE by giving more weight to the fit; but, in contrast to *λ*, this is not by itself sufficient to yield a small SSE (Fig. 3C). However, for small or intermediate *λ*, the differences of SSE and *p*^*⋆*^ between the worst- and the best-ranking paths tend to vanish when *µ* becomes too small (Fig. 3D).

The highest differences between the scores of the best and the worst paths are for *λ* = 10^*−*1.5^ and *µ ≥* 10^*−*2^ (Fig. 3 and Fig. 3D) suggesting that this combination of penalty coefficients could be a good choice for our primary objective of identifying the PIC assembly path from the data. Notably, the best-ranking path is remarkably robust with respect to the choice of the penalty strengths, as soon as *λ* is not too high. In practice, the best-ranking path is the same for all grid points satisfying *λ ≤* 10^*−*1^ for modelT_0_ (Fig. 3E). The heatmaps represented in Supplementary Figs. S3 and S4 still show some variations with *λ* and *µ* in the ranking and relative scores of the suboptimal paths.

### 3.2 Choice of penalty strengths and validation on simulated datasets

To assess the performance of the penalized optimization scheme to identify the PIC assembly paths, and ideally also the steps of PIC assembly altered in each mutant, we simulated data. The simulations encompassed 7 scenarios for the kinetic rates of the wild-type (Supplementary Fig. S5) corresponding to very different occupancy rates for the 6 PIC components (Supplementary Fig. S6). Each of these 7 scenarios was combined with 3 scenarios for which PIC assembly steps are impacted in the two mutants and, for each of these 3 scenarios, the number of altered PIC assembly steps could further be one (Supplementary Fig. S7) or two; in the latter case, the impact of the mutation on the equilibrium constant could be negative for both steps (Supplementary Fig. S8) or negative for one step and positive for the other step (Supplementary Fig. S9). This led to a total of 63 (7x3x3) simulated datasets simulated under model T_0_ on which we conducted parameter optimization under a vast range of penalty strengths (grid of *µ* and *λ* from 10^*−*4^ to 1, Fig. 4A). The results demonstrate the good performance at identifying the PIC assembly path that served to simulate the data, as soon as *λ* is not too high, with relatively similar performance for any *λ ≤*10^*−*1.5^ and no clear impact of the choice of *µ* (Fig. 4B).

**Figure 4:**
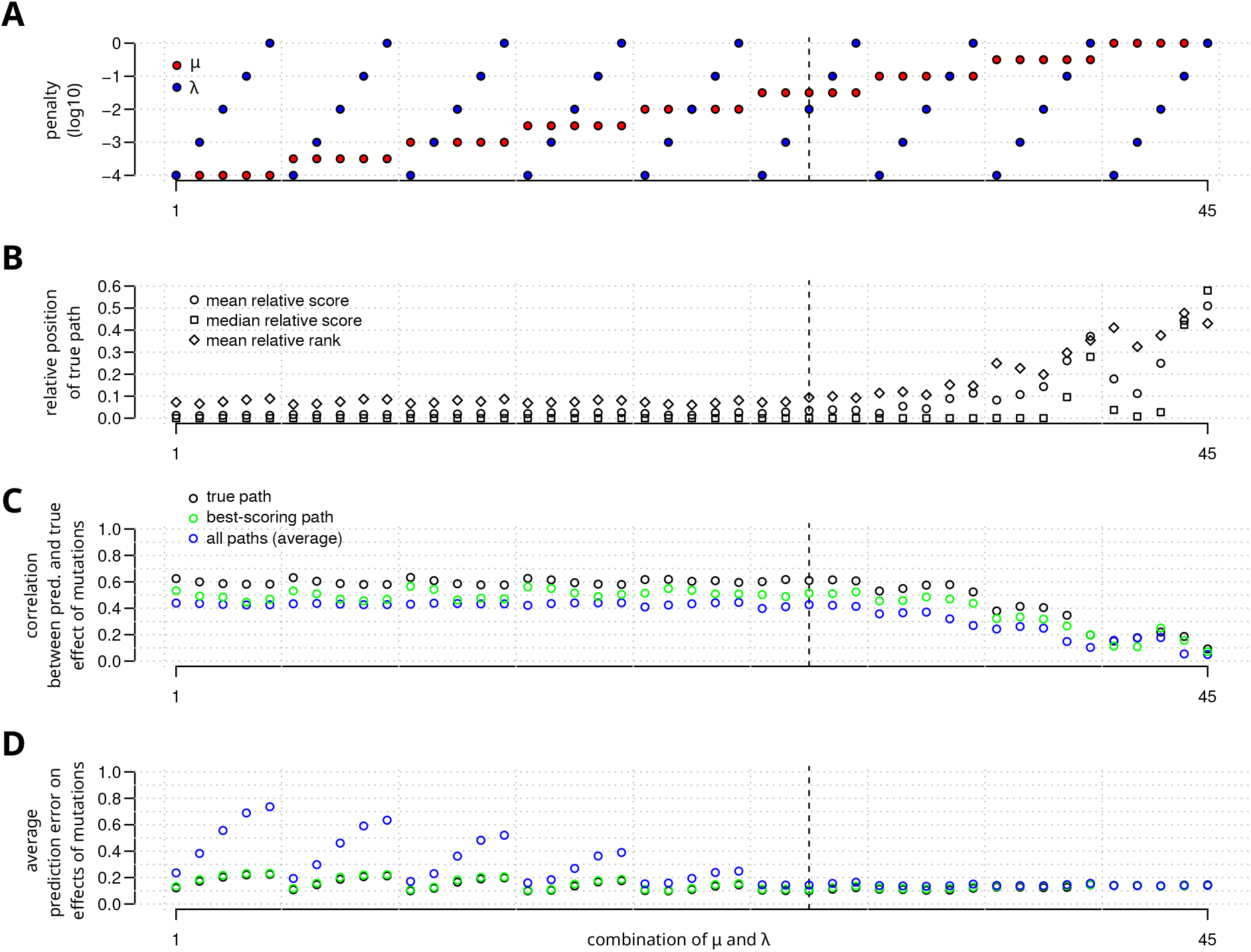
Varying penalty strengths on simulated data. Results obtained for 45 combinations of *µ* and *λ* values (index of combination on x-axis) with transcription model T_0_ on 63 datasets simulated under model T_0_. **A** Values of penalty coefficients *µ* (blue points) and *λ* (red points). **B** Performance of the penalized optimization procedure at predicting the true path. The three types of points show the mean relative score (circles), the median relative score (squares), and the mean relative rank (diamonds) of the true path across the 63 simulated datasets. Both relative scores and relative ranks range from 0 if the true path is the best-scoring path, to 1, if it is the worst-scoring path: relative score is proportional to the score; relative rank is proportional to the number of paths that score better. **C** Pairwise Pearson correlation between the predicted and true impact of the mutations on the recruitment of the 6 PIC components. The impact of a mutation is summarized by the logarithm of its effect on the equilibrium constant of the assembly step corresponding to the recruitment of the component in the path considered 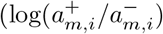, where *m* is the mutant and *i* is the step). Since predictions differ across the 48 possible paths, three values are reported: for the best-scoring path, the true path, and the correlation averaged over all paths. **D** Average absolute difference between the predicted and the true value of the impact of a mutation on a particular assembly step, for the best-scoring path, the true path, and averaged over all paths. The dashed vertical line in **A-D** corresponds to the position of the combination *λ* = 10^*−*1.5^ and *µ* = 10^*−*2^ that we selected for the inference.

In light of these results, we selected for the rest of the analysis *λ* = 10^*−*1.5^ and *µ* = 10^*−*2^. With this penalty strength, the true path was the best ranking path (allowing ties) for 51% (34/63) of the simulated profiles (Supplementary Fig. S10), with an average rank of 4.5/48 (0.094, Fig. 4B). To further use the value of the penalized optimization criterion as an indicator of the support of the data for a path, we recorded the relative position of the value for the criterion within the range between the best and the worst scoring paths. The average relative position of the true path in terms of the optimized criterion was 0.035, which is clearly below the relative position of 0.094 in terms of the sole rank. The relative position therefore provides a more reliable indication of support for a given path than the sole rank because it can better handle cases in which several paths score closely.

Beyond identification of the true path, we also examined the ability to predict which steps of the PIC assembly are altered by the mutations. This ability is confirmed to some extent by the positive correlation between the true and predicted alteration coefficients. Averaged on the 63 simulated scenarios, the Pearson correlation coefficient is 0.51 for the best ranking path and 0.61 for the true path (Fig. 4C). In absolute value the average error on the alteration coefficient of a step is around 0.1 (Fig. 4D) which is relatively high compared to the true value of the alteration coefficients used in the simulation of 0.5 for one or two steps by mutant. This average aggregates diverse types of errors, from small variations around the true values with a globally correct profile to scenarios where the PIC component whose integration is the most impacted are not correctly identified. The results are presented in detail in Supplementary Figures S7, S8, and S9. They allow to better characterize these mistakes which can typically occur when the mutant vs. wild-type occupancy ratios are close for two adjacent PIC state along the assembly path and these components are not ordered properly (see for instance the predictions made for mutant 1 in some simulation scenarios presented in Supplementary Fig. S7B) but also, even when the true path is correctly identified, if the equilibrium constants in the wild-type are strongly biased towards assembly or disassembly (see for instance the predictions made with the scenario 2 for the kinetic rates in Supplementary Fig. S7 and S8). Furthermore, missing data for the last PIC component in our simulation (introduced to mimic the lack of data on TFIIB for *med17-504*) unsurprisingly hampered the identification of the impact of the mutation on the corresponding step.

For the selected penalty strength (*λ* = 10^*−*1.5^ and *µ* = 10^*−*2^), we then used additional simulations to examine the robustness of the inference with respect to different aspects of the data and to the transcription model used for optimization. This included: the level of heterogeneity between kinetic rates governing PIC assembly in the wild-type ; the amplitude of their alteration coefficients in mutants ; the number of PIC assembly steps altered in mutants and the sign of the alteration. We also simulated data with the four different versions of the model for the effect of transcription initiation on PIC assembly and examined the consequences of an inadequacy between the version of the model used for optimization and the version of the model that generated the data. This analysis involved a total of 402 simulated datasets and 810 separate optimizations whose results are summarized in Fig. 5 with all details provided in Supplementary Table S4. For 97.9% of the optimizations the relative score of the true path was below 0.2 (Fig. 5A). The optimizations where it was above could be classified into two types of situations: all paths receive almost the same score, in which case the relative becomes arguably irrelevant; the model used for the optimization assumed a negligible effect of transcription on PIC assembly ( T_0_) whereas the data was simulated with a model in which the transcription could have a significant impact ( T_1_, T_2_, or T_1+2_). These two situations accounted respectively for 7/17 and 10/17 of the 17/810 cases where the relative score of the true path was above 0.2 (Fig. 5B). In this context, it is interesting to note that comparison between the best score obtained under a model with and without transcription could give an indication on the type of model used for the simulation, as seen when examining the 168 datasets were subjected to parameter optimization with several transcription models (Supplementary Fig. S11).

**Figure 5:**
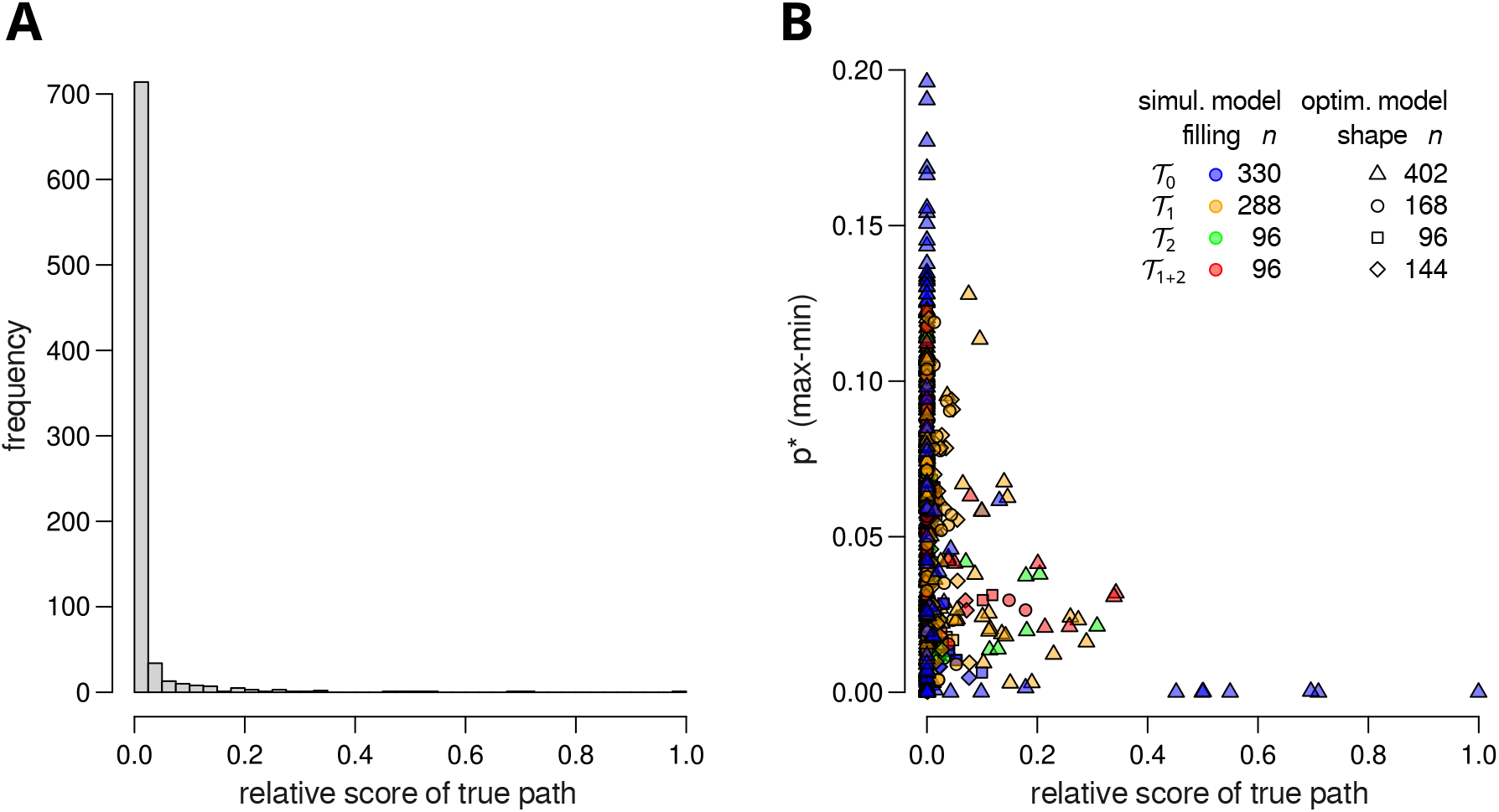
Simulation study to examine the performance of the inference across different transcription models. Results obtained across 810 separate optimizations (covering 402 simulated data sets) with the selected penalty strengths (*λ* = 10^*−*1.5^ and *µ* = 10^*−*2^). **A** Histogram representation of the distribution of the relative score of the PIC assembly path that served to simulate the data (true path). **B** Scatter-plot representation of the relationship between the relative score of the true path and the difference of scores between the worst- and the best-scoring path. Each point represents the result of a separate optimization. For each point, the transcription models ( T_0_, T _1_, T_2_, or T_1+2_) used for the simulation and the optimization aare indicated by the filling color and the shape of the symbol, respectively. The number of points corresponding to each model is reported between parentheses in the inserted legend.

Limitations of the approach pointed by the results on simulated data are also worth mentioning. In models other than T_0_, where transcription initiation introduce non reversibility of the dynamic and thus a distinction between altering the assembly or dissambly rate, the approach proved unable to identifying which of these two rates is altered, in practice the alteration was always reported on the assembly step (not shown). The algorithm proved also unable to estimate the equilibrium constants of the assembly steps in the wild-type (Supplementary Fig. S10CD).

### 3.3 Analysis of profiles corresponding to different groups of genes

For the profile of mutant vs. wild-type occupancy ratios that aggregates the entire gene set, the best-scoring path is stable across a wide range of penalty strengths, from the model T_0_ to T_1+2_ (Fig. 3E, Supplementary Fig. S1E). Details on the score of the whole set of paths, the comparison between predicted and observed profiles, and the associated alteration coefficients are presented in Fig. 6 for model T_0_. A total of four paths received a relative score below 0.2 under model T_0_ (Fig. 6A), and this number rises to six under model T_1+2_ (Fig. 7). The best-scoring path (number 25) corresponds to the recruitment of the 6 PIC components considered in this study in this order: Med, TBP, TFIIK, Pol II, core TFIIH, TFIIB. Taking into account all the paths with a score below 0.2 in T_0_ and T_1+2_, Med is recruited before TBP in all paths but the fourth-ranking path in T_0_. Under model T_0_, path 25 achieves a good fit to the profile. In contrast, the fit of the best-scoring path starting with TBP (path 1) is substantially lower, since it does not capture the fact that Med has a substantially higher mutant vs. wild-type occupancy ratio than TBP in *med17-504* (Fig. 6B).

**Figure 6:**
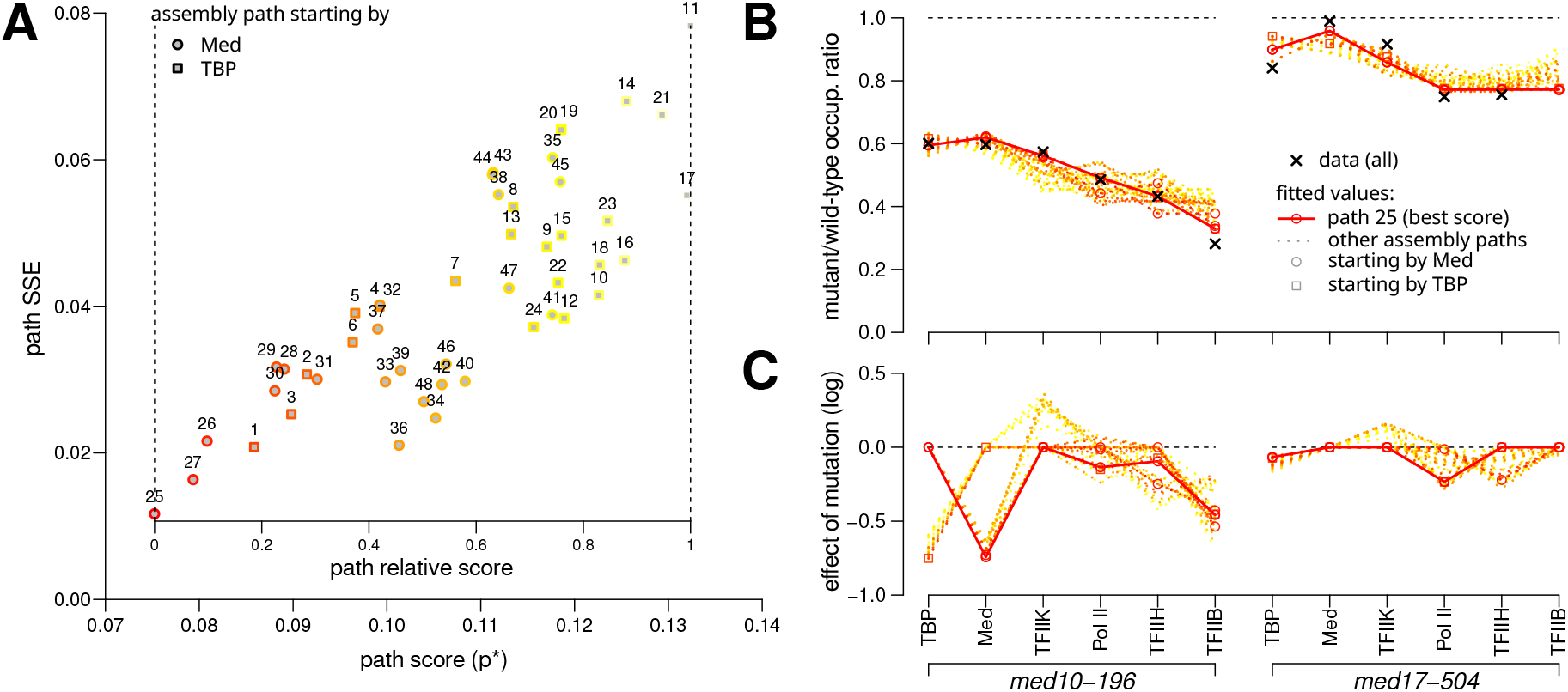
Path scores, goodness of fit and estimated parameters. Results obtained for the entire gene set (dataset “all”) with model T_0_, penalization coefficients *λ* = 10^*−*1.5^ and *µ* = 10^*−*2^. The color reflects the relative score of a path, from red for the best-scoring paths to yellow for the worst-scoring paths. When symbols are used, paths starting with Mediator and TBP are indicated by circles and squares, respectively. **A** Sum of squared errors (SSE) of prediction versus score (*p*^*⋆*^) for the 48 evaluated PIC assembly paths. **B** Comparison between predicted and observed mutant/wild-type occupancy profiles associated with the 48 paths. The best-scoring path, which is here also the best path in terms of SSE, is represented by a plain line and other paths by dotted lines. Paths with relative score *≤* 0.2 are also highlighted by symbols. The experimental data on which the model was fitted (dataset “all”) are represented by black crosses. **C** Predicted impact of the alteration caused by the mutations on the recruitment of each PIC component 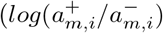, where *m* is the mutant and *i* the step). Colors and symbols for the paths are as in **B**.

**Figure 7:**
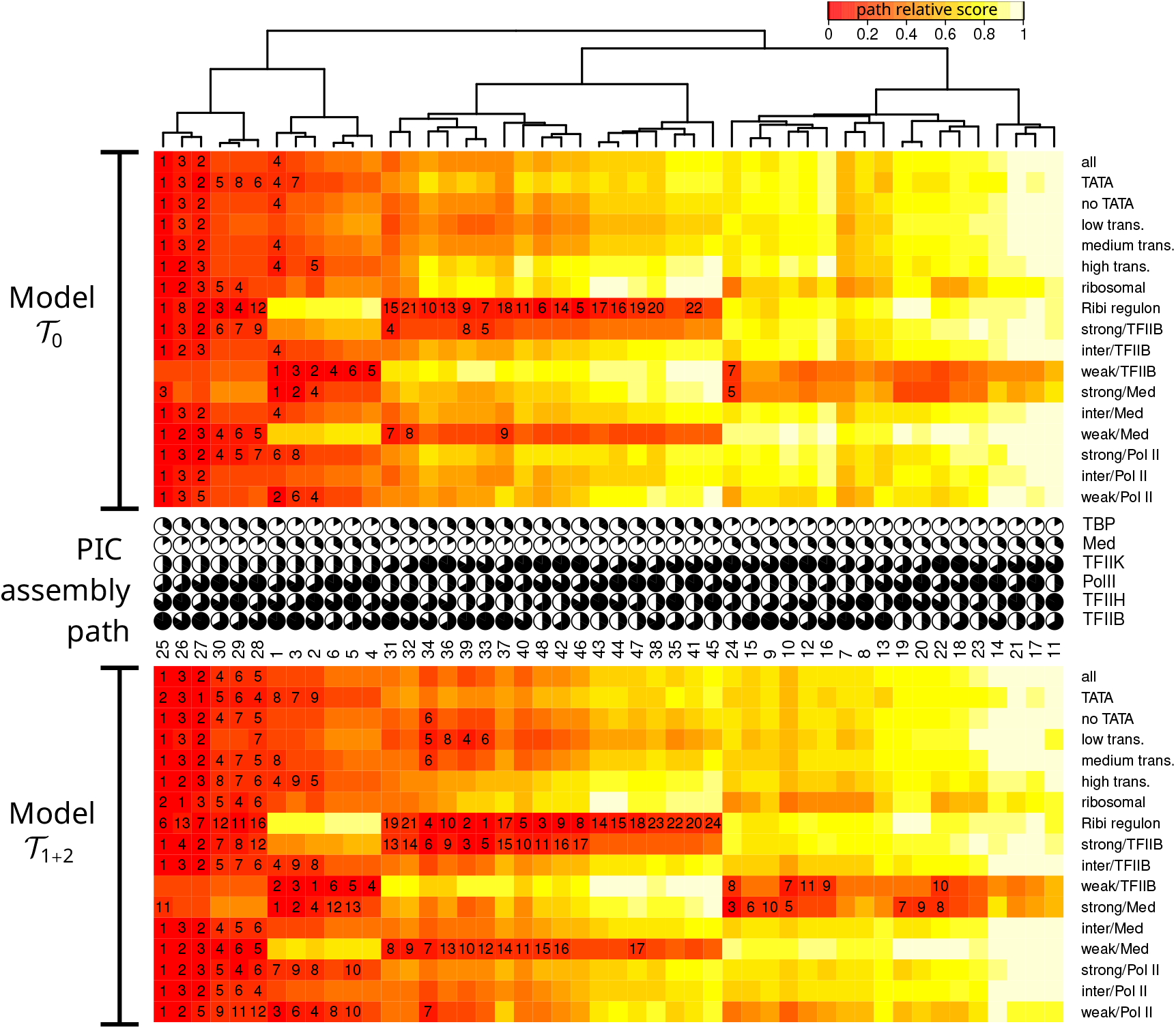
PIC assembly paths clustered by ranking on 17 groups of genes. Upper heatmap: model that neglect the effect of transcription initiation on PIC assembly (model T_0_, reversible). Lower heatmap: model with transcription and reinitiation at empty promoter and PIC assembly step before Pol II entry (model T _1+2_). Penalization coefficients: *λ* = 10^*−*1.5^, *µ* = 10^*−*2^. Each cell corresponds to one of the 48 PIC assembly paths compared in this study (columns) and one of the 17 data sets (rows). The color reflects the relative score of the path for the considered dataset and model of transcription. Numbers printed within heatmap cells indicate the rank of paths with a optimization criterion relative position below 0.2. The top tree was built by average-link hierachical clustering on pairwise distances between the vectors of optimization criterion relative positions across datasets and models ( T_0_, T_1_, T_2_, and T_1+2_).Circular diagrams with 6 sectors filled clockwise indicate the recruitment steps of the PIC components in the assembly paths.

Among the paths for the entire gene set, a common feature to all those scoring below 0.2 is the recruitment of TFIIK third, before the core module of TFIIH (Fig. 7). This is consistent with previous studies of yeast Mediator mutants that suggested independent recruitment of TFIIH’s two modules [7, 15, 33], and is interesting, since the ordered PIC assembly inferred *in vitro* did not distinguish these two modules within TFIIH. For comparison, the path inferred *in vitro* corresponds to the recruitment of the PIC components in this order: TBP, TFIIB, Pol II, TFIIH. This would correspond to paths 45 and 46 when Med is recruited before TBP, and to paths 21 and 22 when TBP is recruited before Med.

On this entire gene set, three main predicted alterations of PIC assembly steps stand out in the prediction of the effects of the mutations. They concern, in order of amplitude, the recruitment of Med itself (or TBP for the path in which it arrives first) and TFIIB, both for the mutant *med10-196*, and that of Pol II, for the mutant *med17-504* (Fig. 6C). These estimations are consistent with the previously identified impacts of these mutations on physical interactions of Mediator with TFIIB or Pol II, respectively [7, 15]. In particular, we previously showed that the Mediator mutant *med10-196* led to a decrease in the physical interaction of Mediator with TFIIB [7].

In [34], the binding dynamics of several PIC components were investigated by competition ChIP in *S. cerevisiae*, and this work revealed that PIC dynamics depend on the gene, and showed a longer residence time for ribosomal genes or gene promoters with strong TATA-elements. However, it remains to be investigated how assembly paths differ between groups of genes. To refine our analysis in this direction, we analyzed the mutant vs. wild-type occupancy profiles of 16 different subgroups of genes, beside the entire gene set. Among these, 3 groups reflect the transcriptional activity, measured by Pol II occupancy of the corresponding genes in the wild-type context, and 2 groups correspond to the presence or absence of a TATA box in the gene promoter. We also included the groups of ribosomal protein genes and the ribosome biogenesis (Ribi) regulon [29], that have specific transcription factors and a coordinated mode of transcription regulation. A total of 9 other groups of genes were defined according to the effect of the Mediator *med10-196* mutation on Pol II, TFIIB, and Mediator binding, and were also considered, in keeping with [7]. We first compared, on these 17 profiles, the fit of the four different transcription models (Fig. 7). We observe that the number of paths with a relative score below 0.2 tends to increase with the number of parameters in the transcription model, but, in general, the best-scoring paths were comparable between model T_0_ and model T_1+2_. When comparing the models in terms of the raw score *p*^*⋆*^, which we had seen could be indicative of the model that generated the data in our simulated study, we found that the fit is better with a model that accounts for transcription only for four subgroups of genes, namely strong/Med, weak/TFIIB, Ribi regulon, and ribosomal (Supplementary Fig. S12). This overall low power of the modeling of transcription initiation to improve the fit might well reflect the fact that completing PIC assembly does not necessarily result in productive transcription. In human cells, it has been proposed that only a small fraction of PIC assemblies result in elongation, and that the turnover of the Pol II initiation complex is high compared to the transcription initiation rate [35, 36].

Major differences in path support, compared to the entire gene set, were observed for 3 out of 16 subgroups of genes. These subgroups represented two different signatures in terms of the distribution of path supports across the 48 paths considered (Fig. 7, Supplementary Fig. S13). The first signature is shared by the strong/Med and weak/TFIIB groups (Supplementary Fig. S13AB), whose best-scoring paths start with TBP instead of Med, as observed for the entire gene set. The second signature is found in the Ribi regulon group, (Supplementary Fig. S13C) whose best-scoring paths start with Med but lack the clear signal of TFIIK in third position found for the entire gene set. These results suggest that PIC assembly paths could depend on the group of genes, potentially constituting an additional level of regulation. It is striking that these three subgroups are among those for which the models with transcription achieve a better fit than the model without transcription (Supplementary Fig. S12). With respect to the different positions of TBP in the paths associated with some subgroups of genes, it is interesting to note that TBP dynamics could be one of the critical elements contributing to the differences between groups of genes. For example, Rap1-dependent genes were shown to have slower TBP turnover compared to other Pol II-transcribed genes [18, 20].

A final point of notice is that the fit is generally not as good on the experimental data as on the simulated data. This is particularly true for the strong/Med and Ribi regulon groups, for which the best score is above 0.10 (Supplementary Fig. S12), while it is below this value in all of the 810 simulated datasets (Supplementary Fig. S11). This may, for instance, be due to more complex alterations of the PIC assembly paths caused by the mutations than those taken into account in the model, or to more complex assembly paths that may not be well represented by a single linear sequence of recruitment.

## 4 Conclusion

In this study, we developed a quantitative framework that scores PIC assembly paths according to their ability to explain experimental data consisting of mutant vs. wild-type occupancy profiles of the PIC components. To our knowledge, this is the first modeling approach developed to account for the impact of Mediator mutants on genome-wide occupancy profiles of PIC components measured by ChIP-seq. This work improves our understanding of PIC assembly paths, opening perspectives for future modeling and experimental studies.

The main findings can be summarized in six points: (i) the best-supported PIC assembly paths are different from the sequence inferred *in vitro*; (ii) multiple paths fit the experimental data equally well; (iii) introducing the reinitiation of PIC assembly upon transcription, either to the empty promoter or at a different step of the assembly process, has relatively little impact on which paths are the best-scoring, but increases the number of paths that become compatible with the data; (iv) important differences in the support of assembly paths were observed for specific groups of genes, suggesting that the path could depend on the gene and may constitute a level of regulation of gene expression; (v) paths starting with Mediator or TBP but otherwise similar receive different levels of support, emphasizing the importance of taking into account both components; (vi) the respective positions of the core TFIIH module and the TFIIK kinase module within the best-supported paths are consistent with their behavior as two distinct components.

Our quantitative framework for scoring PIC assembly paths could be further applied to data from other mutants perturbing PIC assembly at different levels, which may reduce the number of supported paths. Future work integrating a more global model and several types of data may also encompass transcription machinery assembly and transcriptional activity. However, this probably requires accounting for complex biological phenomena such as the existence of transcription bursts [37–39]. Also, mammalian Mediator was recently proposed to play a role in controlling transcription reinitiation through the formation of Pol II convoys [40]. Our approach constitutes a step towards more quantitative understanding of PIC assembly within the complex mechanisms of transcription regulation.

## Acknowledgments

This work was supported by the Fondation ARC (grants SL220130607079 and PGA1 RF201702055342) and the Agence Nationale de la Recherche (ANR-14-CE10-0012-01). We thank M. Werner for helpful discussions on PIC assembly.

## Supplementary figures

**Figure S1:**
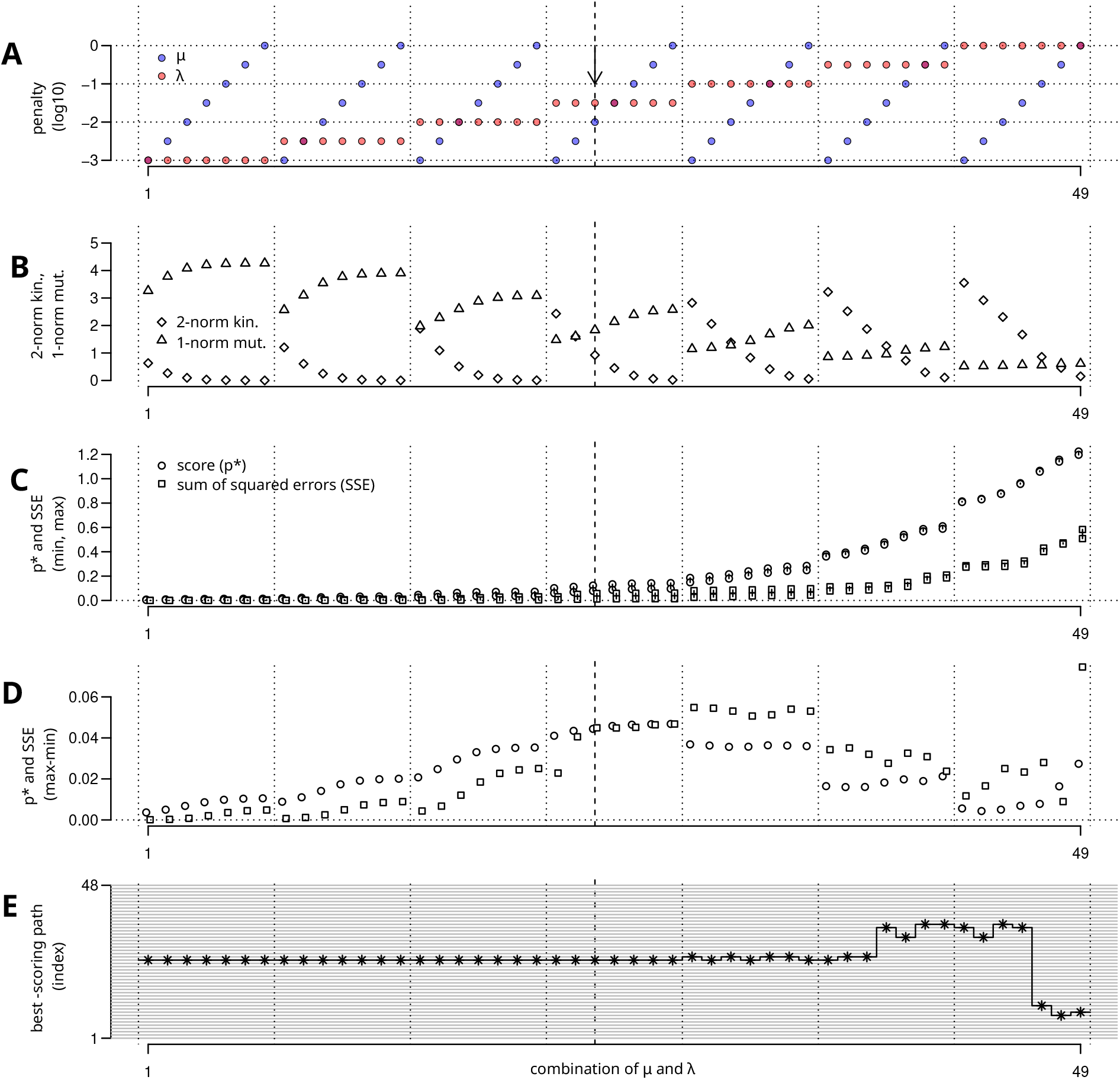
Varying penalty strengths on real data, results with model. T_1+2_. Results obtained on the entire gene set when varying *µ* and *λ*. Same representation as in Main text Figure 3, but for transcription model T_1+2_. See the corresponding legend in main text.

**Figure S2:**
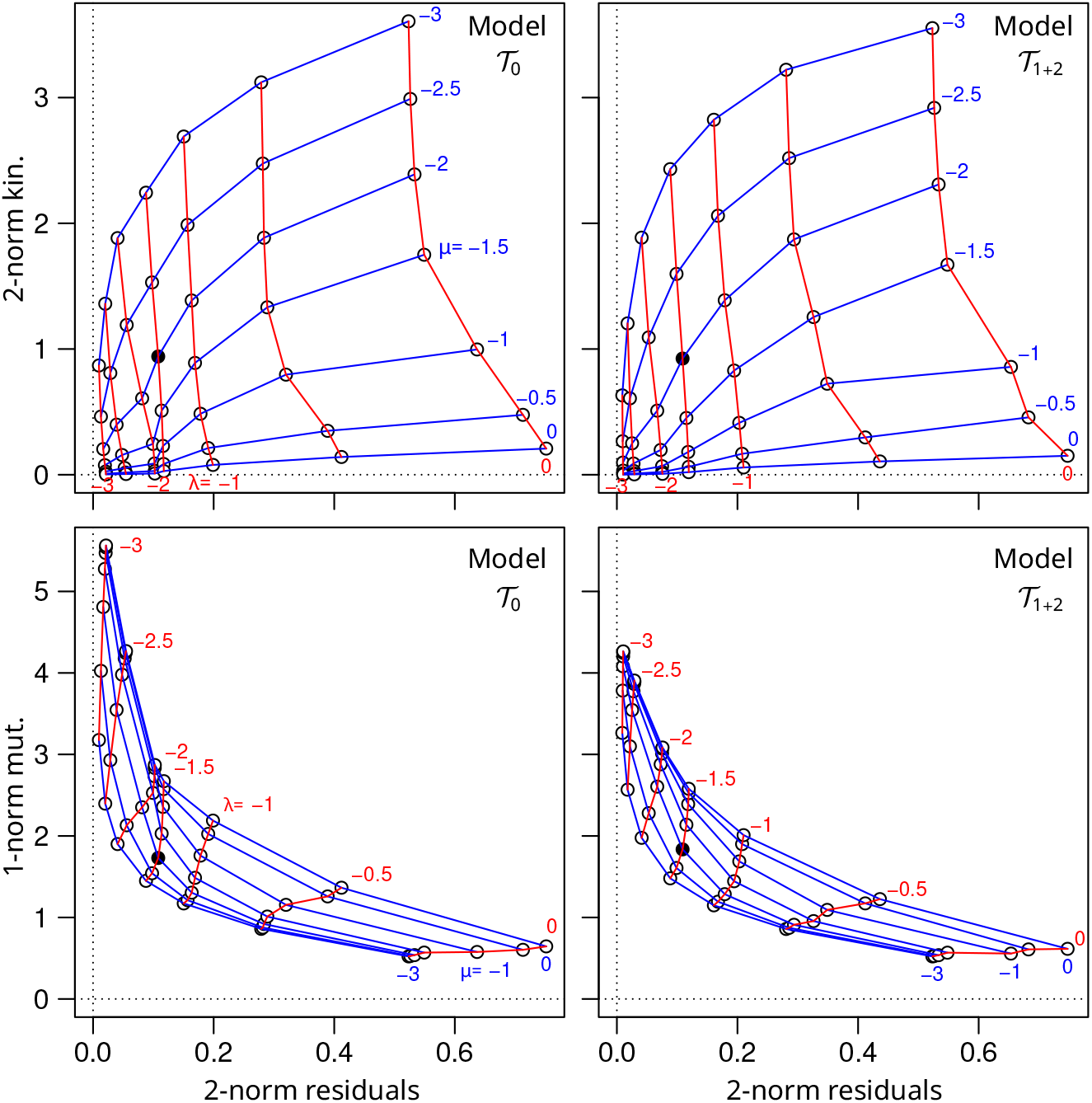
L-curves. Representation based on results obtained on real data (entire gene set) with models T_0_ (left column) and T_1+2_ (right column). The name L-curve refers to the typical shape of these curves that represent the solution norm (y-axis) versus the residual norm (x-axis) for different penalty strengths [30]. The typical L-shape is observed in the lower row but not in the upper row. The x-axis is the residual 2-norm (square root of the SSE) in the four plots. The y-axis differ between the upper and lower rows, showing the 2-norm of kinetic rates ( l*∥*og(*k*) *∥* _2_) and the 1-norm of their alteration coefficients in mutants ( *∥log*(*a*) *∥*_1_), respectively. Each point corresponds to the solution (best parameter values for the best path) obtained for a combination of *µ* and *λ* values (49 combinations). The lines connect the points associated with a same value of *µ* (blue lines) or *λ* (red lines). Some values of *µ* and *λ* are printed in blue and red, respectively. The combination *µ* = 10^*−*2^ and *λ* = 10^*−*1.5^ retained for the inference is shown as a black point.

**Figure S3:**
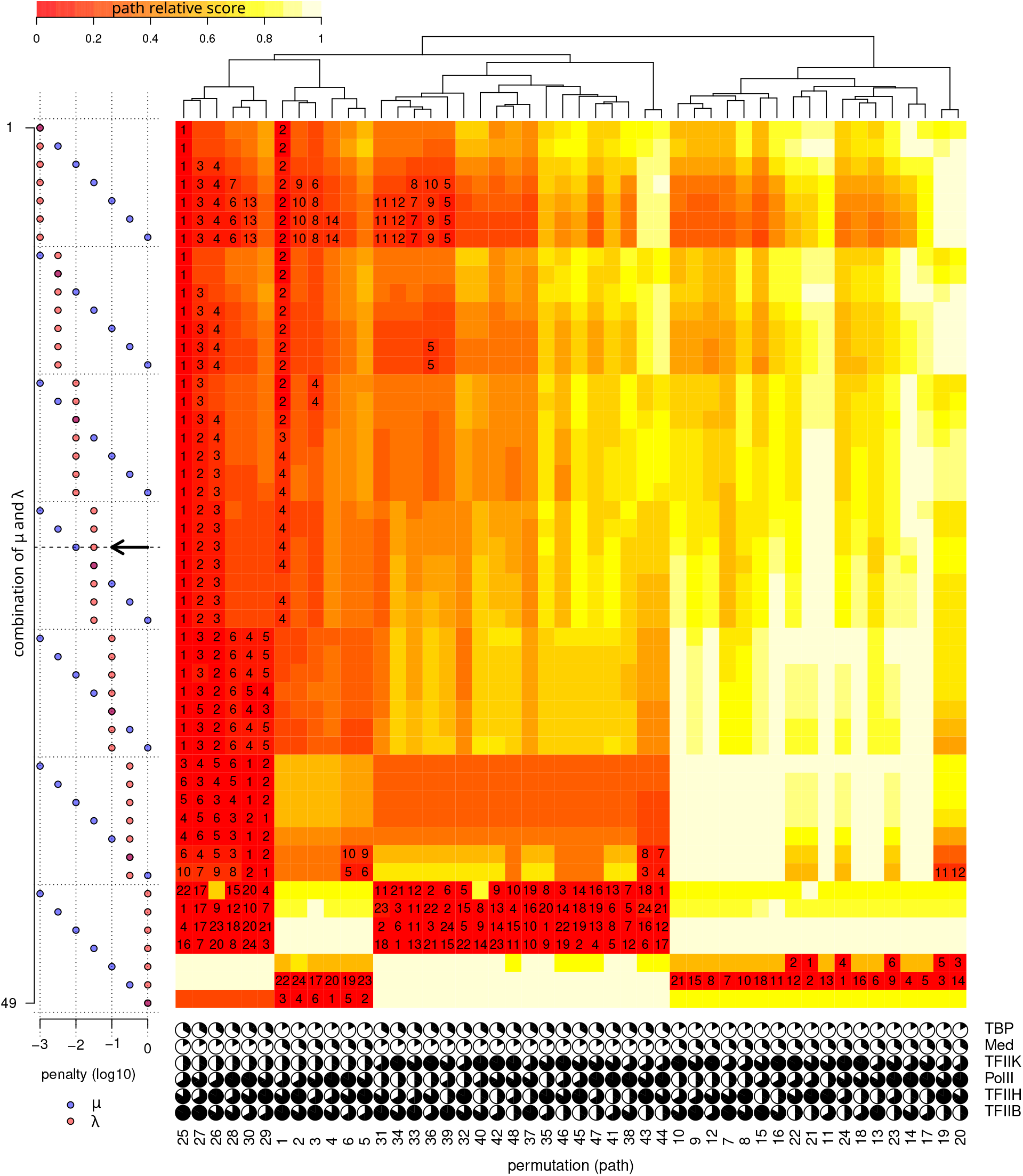
Impact of penalty strengths on path scores, results with model. T_0_. Heatmap representation of the results obtained on the entire gene set with the transcription model T_0_ for 49 combinations of *µ* and *λ* values (rows). Each column corresponds to one of the 48 possible PIC assembly paths. The top tree that served to order the columns was built by average-link hierachical clustering on pairwise distances between the vectors of path relative scores across the *µ* and *λ* values for models T_0_ and T_1+2_ (same tree in Supplementary Figures S3 and S4). The color reflects the path relative score for a combination of *µ* and *λ* values. The values of penalization coefficients *µ* and *λ* are represented on the left side of the heatmap.

**Figure S4:**
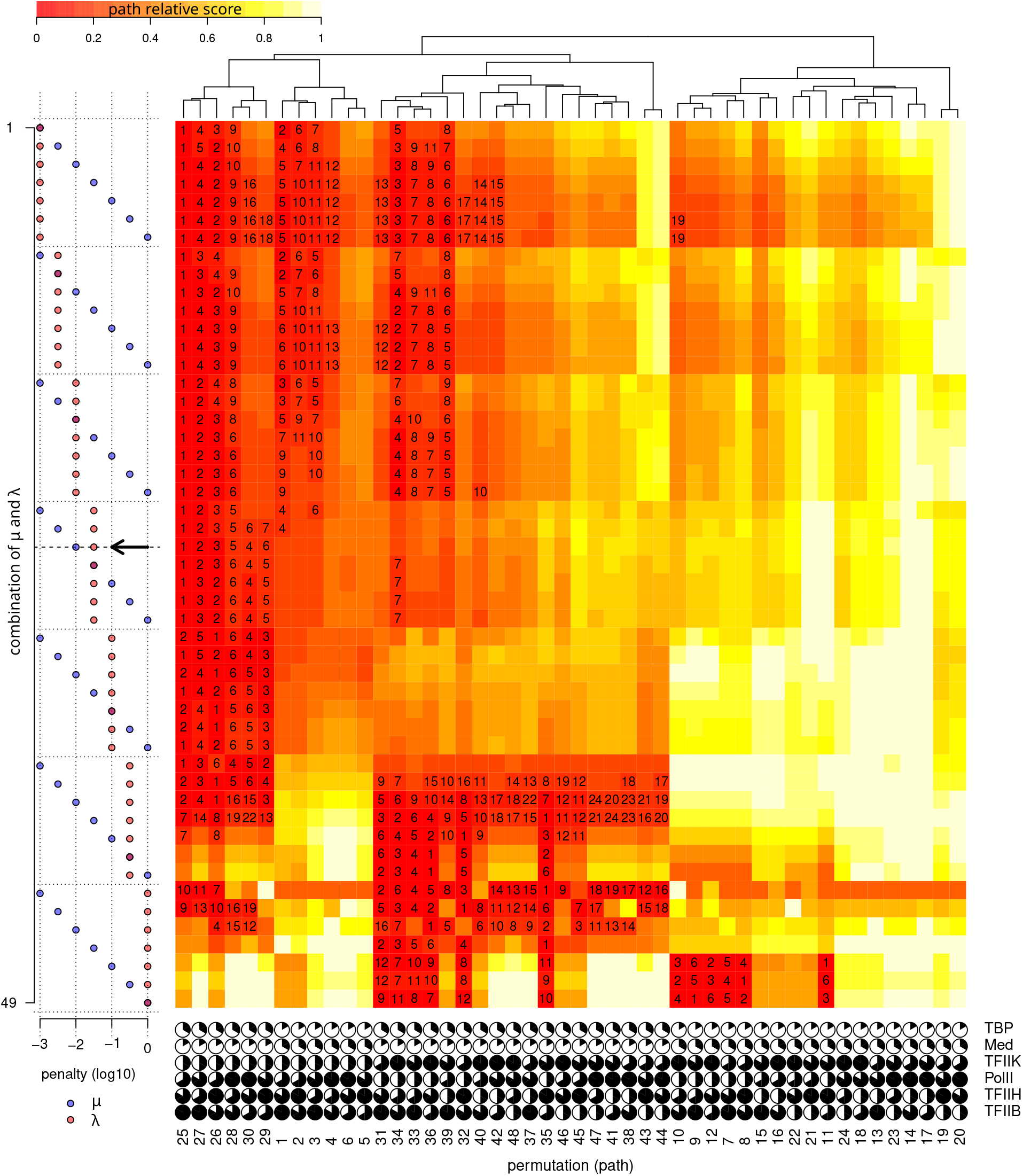
Impact of penalty strengths on path scores, results with model. T_1+2_. Results obtained on the entire gene set when varying *µ* and *λ*. Same representation as in Supplementary Figure S3, but based on results obtained with the model T_1+2_. See the corresponding legend.

**Figure S5:**
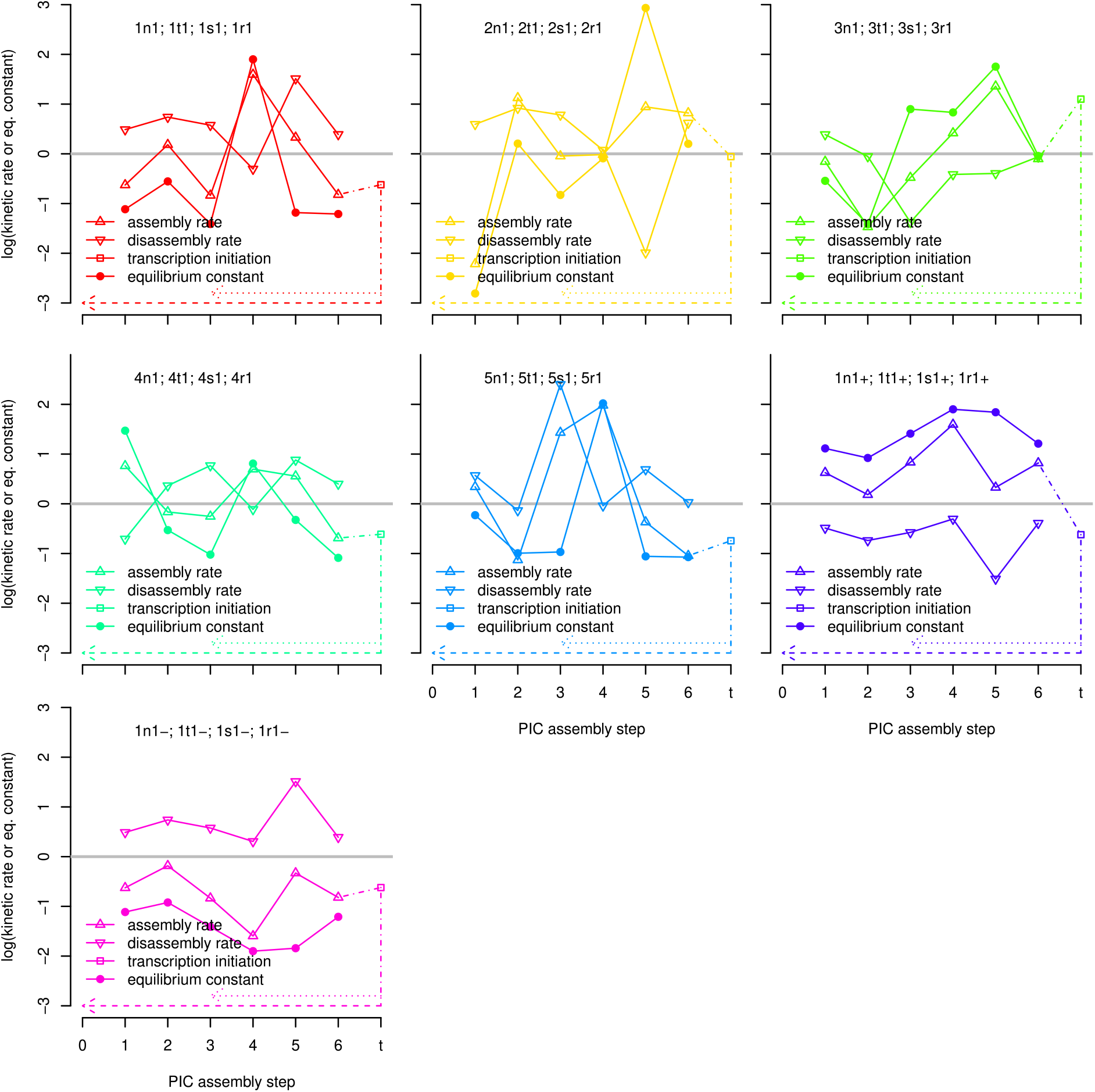
Simulated scenarios for the kinetic rates of the wild-type. The 7 sets of kinetic parameters are shown in different panels and represented using different colors. In each panel, we show the logarithm of the assembly/disassembly kinetic rates 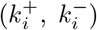 and the equilibrium constant 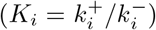 of the 6 PIC assembly steps that determine the model T_0_, as well as the kinetic rate for the transitions corresponding to transcription initiation (*k*^+^) that are added in models T_1_, T_2_, and T_1+2_. Since two transitions are added in T_1+2_, the total transcription rate is twice higher in T_1+2_ than in T_1_ and T_2_. The sets of kinetic rates presented in the 7 panels are respectively labeled 1x1, 2x1, 3x1, 4x1, 5x1, 1x1+, and 1x1-, where x is a one-letter code that indicates the transcription model (“n” for T_0_, “r” for T_1_, “s” T_2_, and “t” for T_1+2_), the first number indicates the set of kinetic rates, the second number indicates the multiplicative coefficient applied to the log of the assembly/disassembly kinetic rates. This multiplicative coefficient tunes the level of heterogeneity between kinetic rates governing PIC assembly in the wild-type (“dispersion factor”, always 1 in this figure but also takes the values 0, 0.5 and 2 in the simulations). The log of the kinetic rates for the first 5 scenarios were randomly drawn from a centered Gaussian distribution with unit variance. The last two sets reuse the values of the first set but exchange the assembly and disassembly rates when needed such that the equilibrium constant is, for all steps, in favor of the assembly for the sixth set and in favor of the disassembly for the seventh set. These two sets are distinguished from the first set by labels that terminates with “+” and “-”.

**Figure S6:**
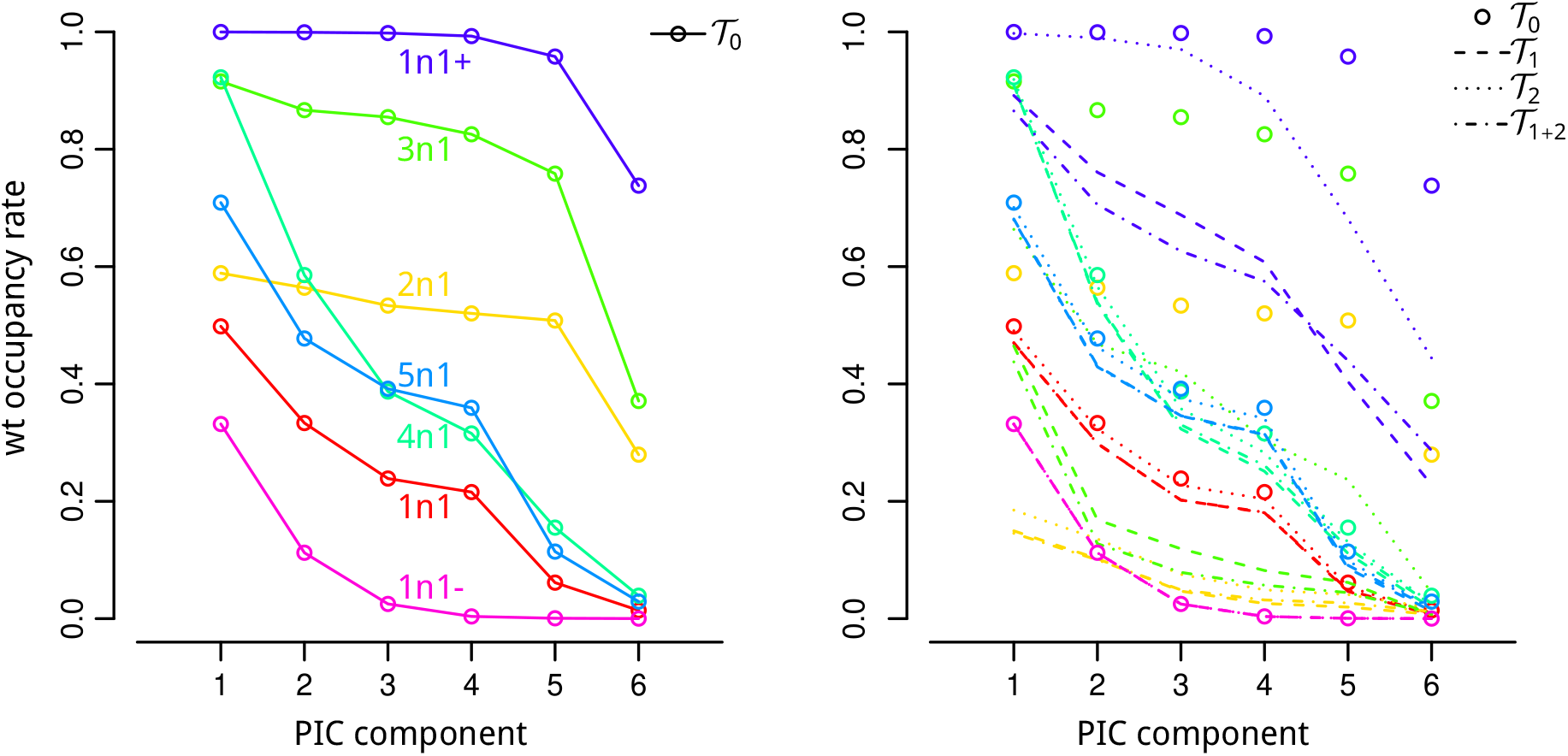
Occupancy rates of the 6 PIC components in wild-type for the simulated scenarios. Left panel: occupancy rates under model T_0_ for the 7 sets of kinetic parameters (distinguished by different colors). The kinetic rates that determine these occupancy profiles are presented in Fig. S5. Right panel: change in occupancy rates when transcription initiation is added. The occupancy profiles under models T_1_ (dashed lines), T_2_ (dotted lines), and T_1+2_ (dotted-dashed lines), are compared to those under model T_0_ (open circles). For instance, the models labeled 1r1, 1s1, and 1t1, are compared to the model labeled 1n1 (red).

**Figure S7:**
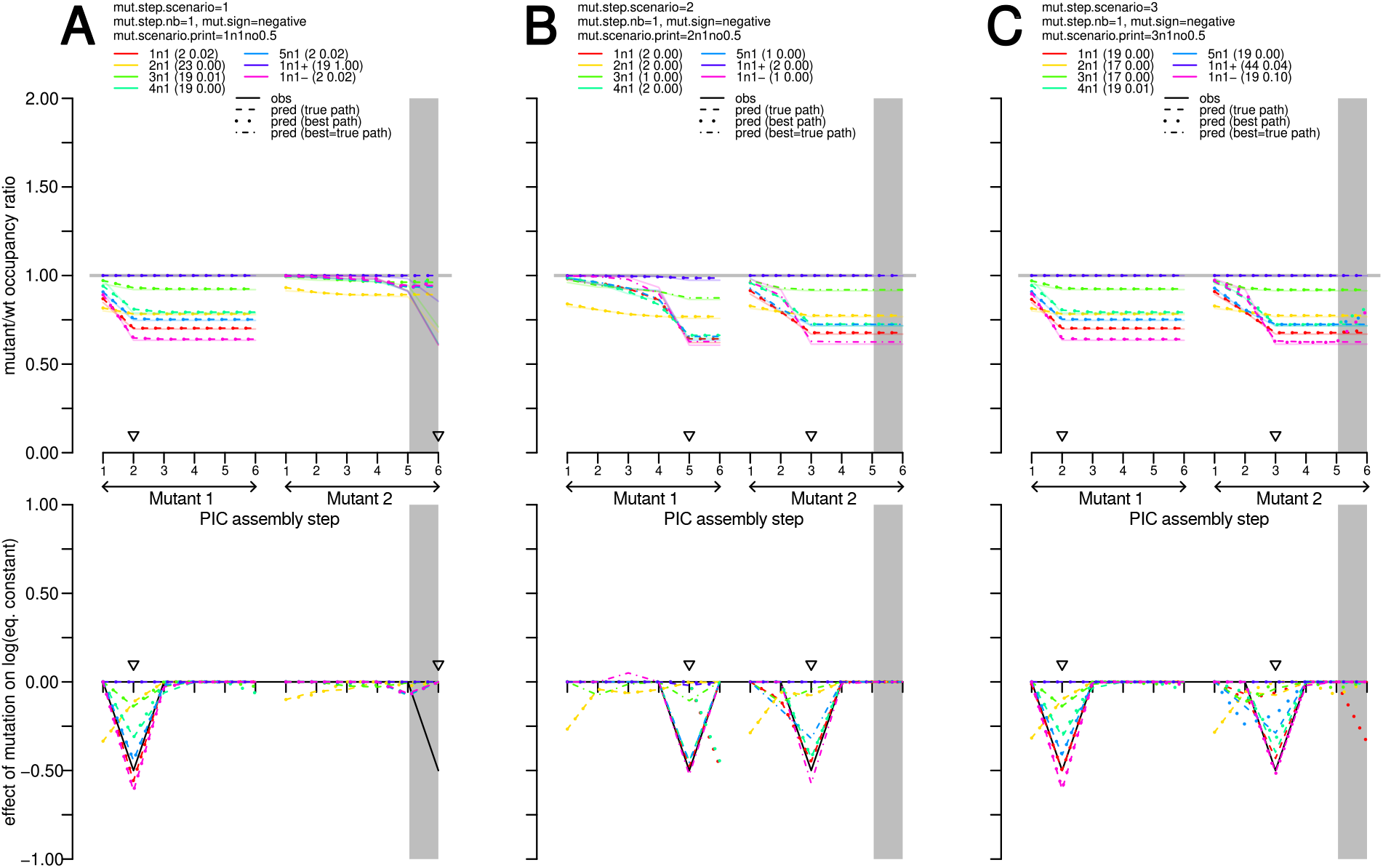
Results on simulated data for different scenarios with a single PIC assembly step negatively impacted in each mutant. The three panels (**A, B, C**) distinguishes the three sets of simulated scenarios for the impact of the mutations. The impacted PIC steps were selected randomly and are indicated by downward triangles along the x-axis. Like the labels attributed to the scenarios for the kinetic parameters of the wild-type (Fig. S5), each scenario for the impact of the mutations is identified by a unique label. This label is for instance “1n1no0.5” for the panel **A**, it is composed of: a number in first position (which can be 1, 2, or 3; here 1) indicating the set to which the scenario belongs; a character in second position (which can be “n” for “negative” or “m” for “mixed”; here “n”) indicating the sign of the alteration of the equilibrium constant; a number in third position (which can be 1, 2, …, 6; here 1), indicating the number of PIC assembly steps altered in each mutant; a two-letter in fourth position (which can be “no”, “ka”, or “kd”, for “not relevant”, “assembly rate”, “disassembly rate”; here “no”), indicating whether the alteration of the equilibrium constant is caused by an impact on the assembly rate, on the disassembly rate, or whether this distinction is irrelevant (case of model T_0_ whose occupancy profiles are determined by the equilibrium constants); a number in last position (which can be 0.25, 0.5, or 1; here 0.5) indicates the amplitude of the alteration coefficients in the mutants (an additive coefficient applied to log of the equilibrium constant). The upper plot of each panel shows the mutant vs. wild-type occupancy ratios that served for the inference. The different colors distinguish the 7 scenarios determined by the 7 sets of kinetic parameters presented in Fig. S5 which are identified by their labels in the insert legend. For each scenario, the rank and relative score of the true path is indicated alongside the label, between parentheses. The lower plot of each panel shows the predicted and true impact of the alteration (black and colored lines, respectively) caused by the mutations on the recruitment of each PIC component 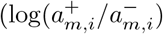, where *m* is the mutant and *i* the step; see Fig. 4). Since the predicted impact depends on the path, it is reported here for the true path (that served to simulate the observed dataset) and for the best-scoring path. Dash and dotted lines are used for the true and the best-scoring paths, respectively. When the best-scoring path is also the true path a dotted-dashed line is used. All the results presented in this figure were obtained with model T_0_ and penalization coefficients *λ* = 10^*−*1.5^ and *µ* = 10^*−*2^.

**Figure S8:**
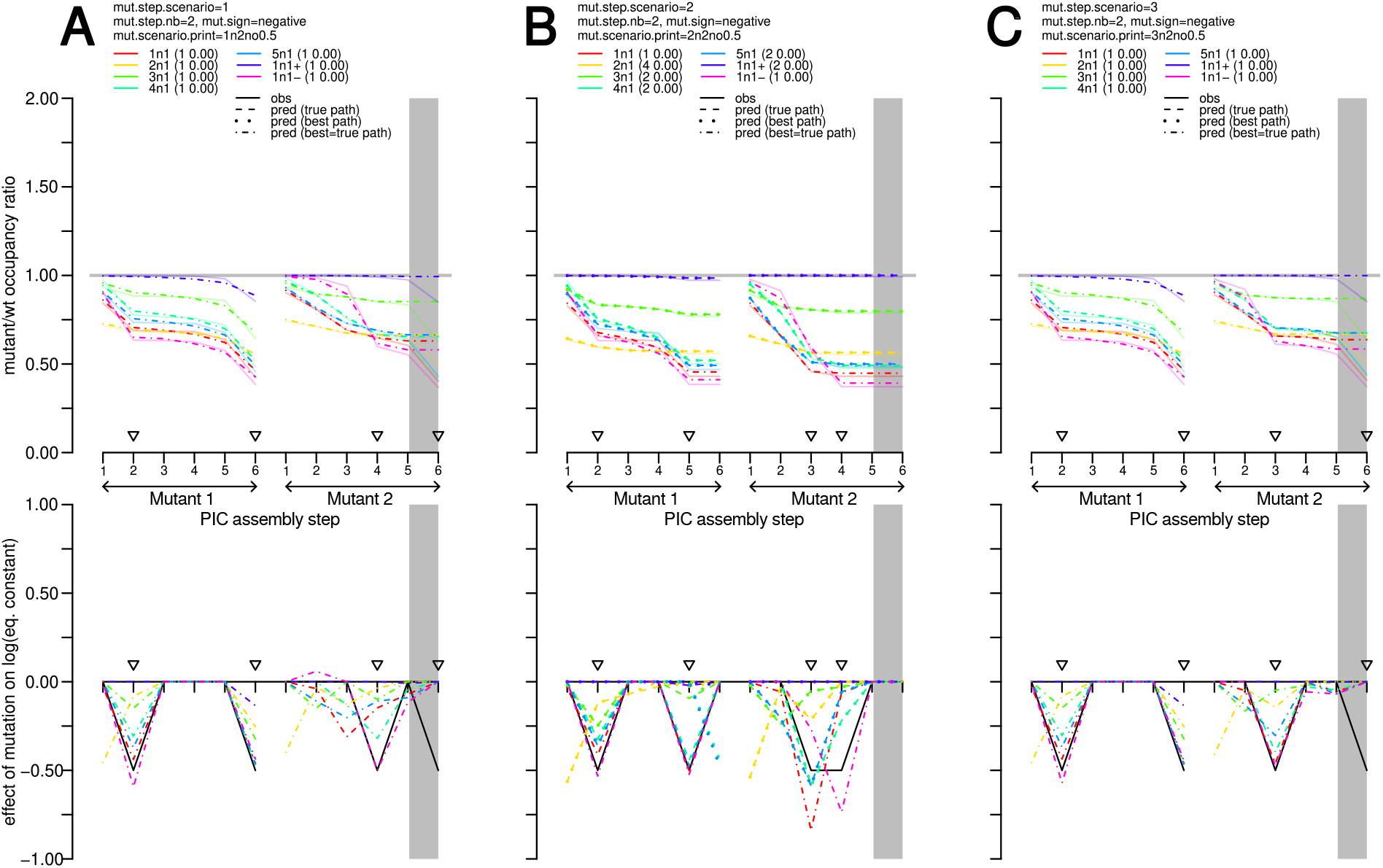
Results on simulated data for different scenarios with two PIC assembly steps negatively impacted in each mutant. See the legend of Fig. S7.

**Figure S9:**
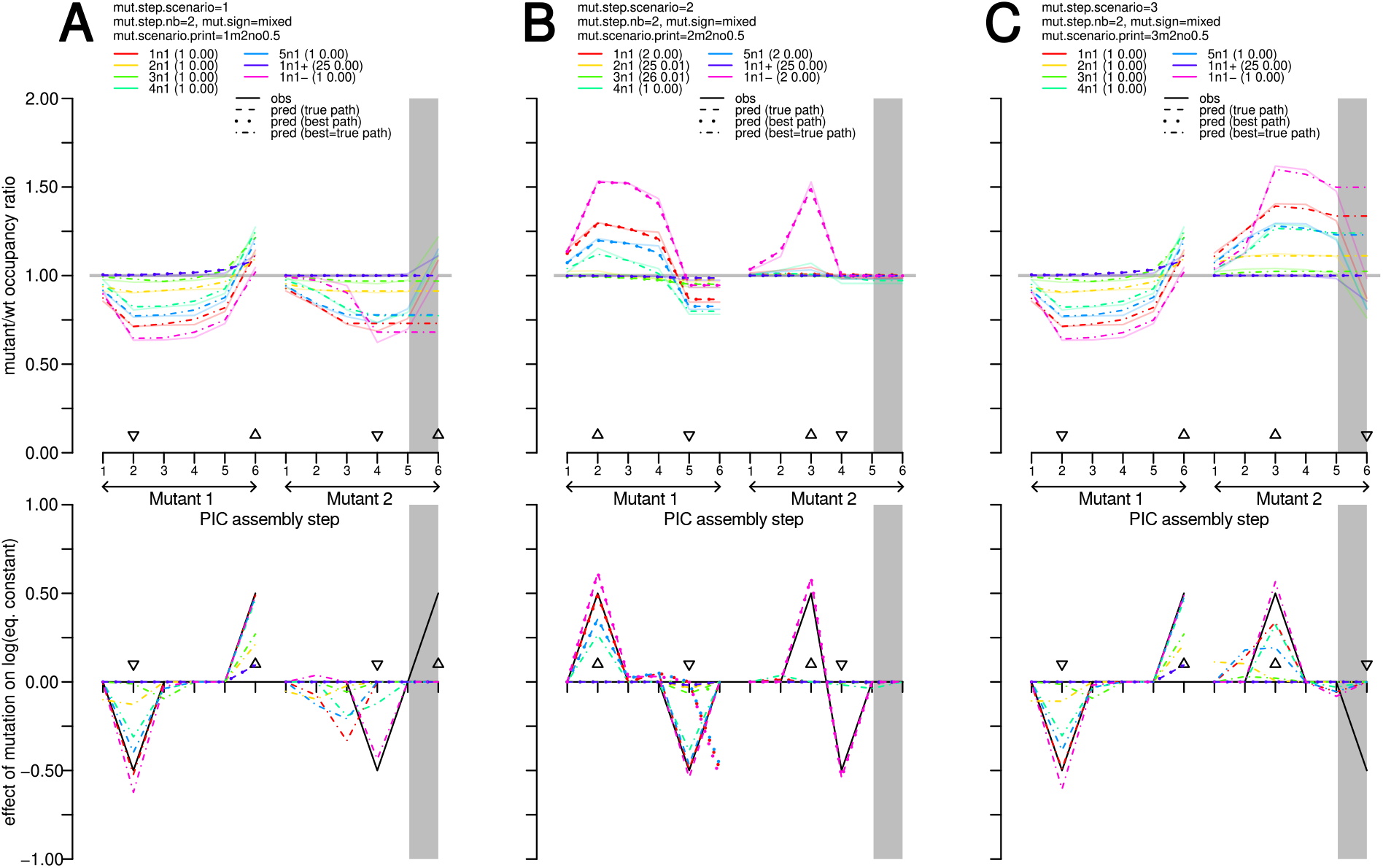
Results on simulated data for different scenarios with two PIC assembly steps impacted in each mutant, negative and positive alteration coefficients. See the legend of Fig. S7.

**Figure S10:**
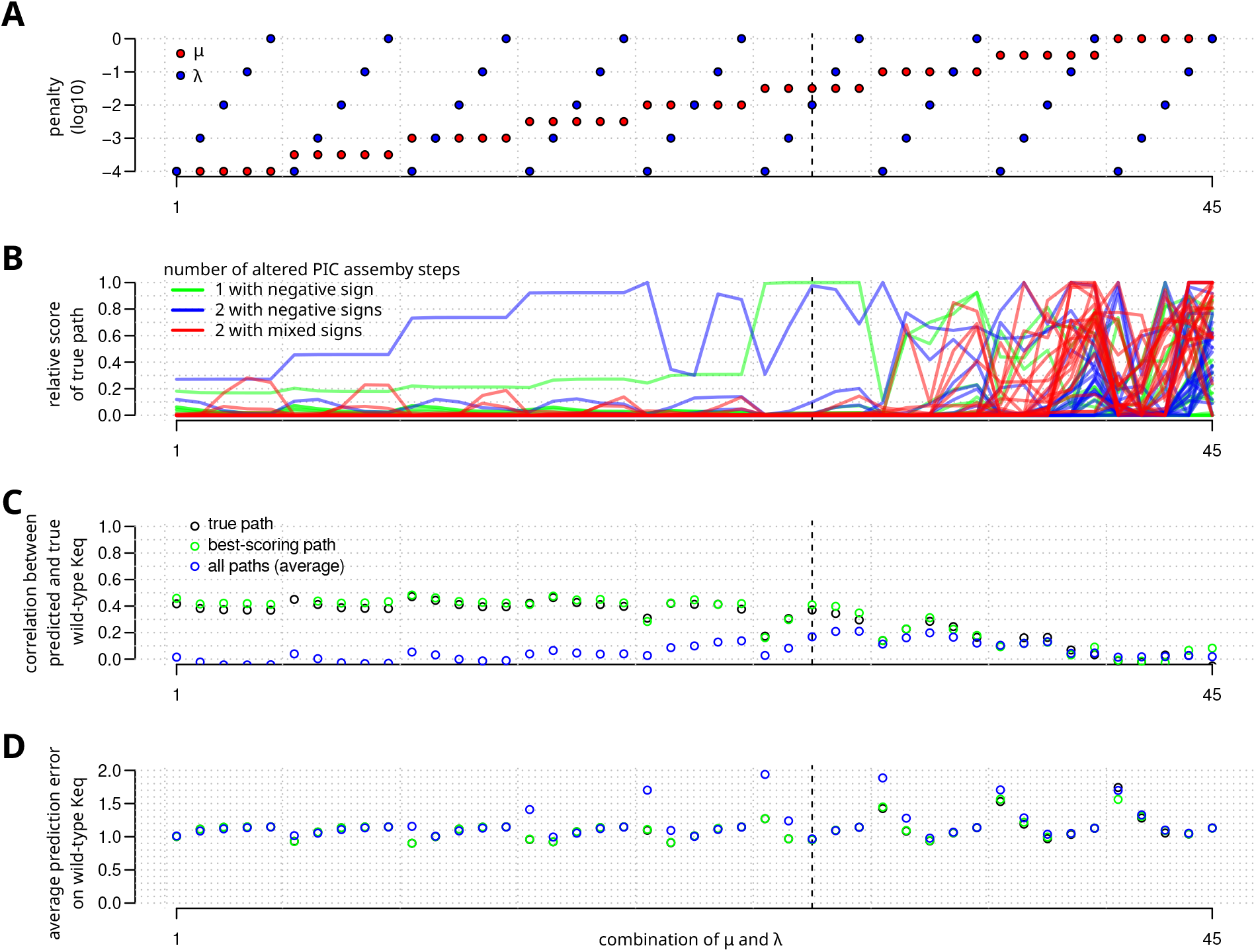
Additional details on results of varying penalty strengths on simulated data. Complementary to main text Fig. 4. **A** Values of penalty coefficients *µ* (blue points) and *λ* (red points). **B** Relative score of the true path across the grid of *µ* and *λ* values for each of the 63 simulated datasets. **C** Correlation between the predicted and true values of the equilibrium constants (log) in the wild-type, averaged on the 63 simulated datasets. Since predictions differ across the 48 possible paths, three values are reported: for the best-scoring path, the true path, and the correlation averaged over all paths. **D** Average absolute difference between the predicted and the true value of the equilibrium constant (log) of a particular assembly step in the wild-type, for the best-scoring path, the true path, and averaged over all paths.

**Figure S11:**
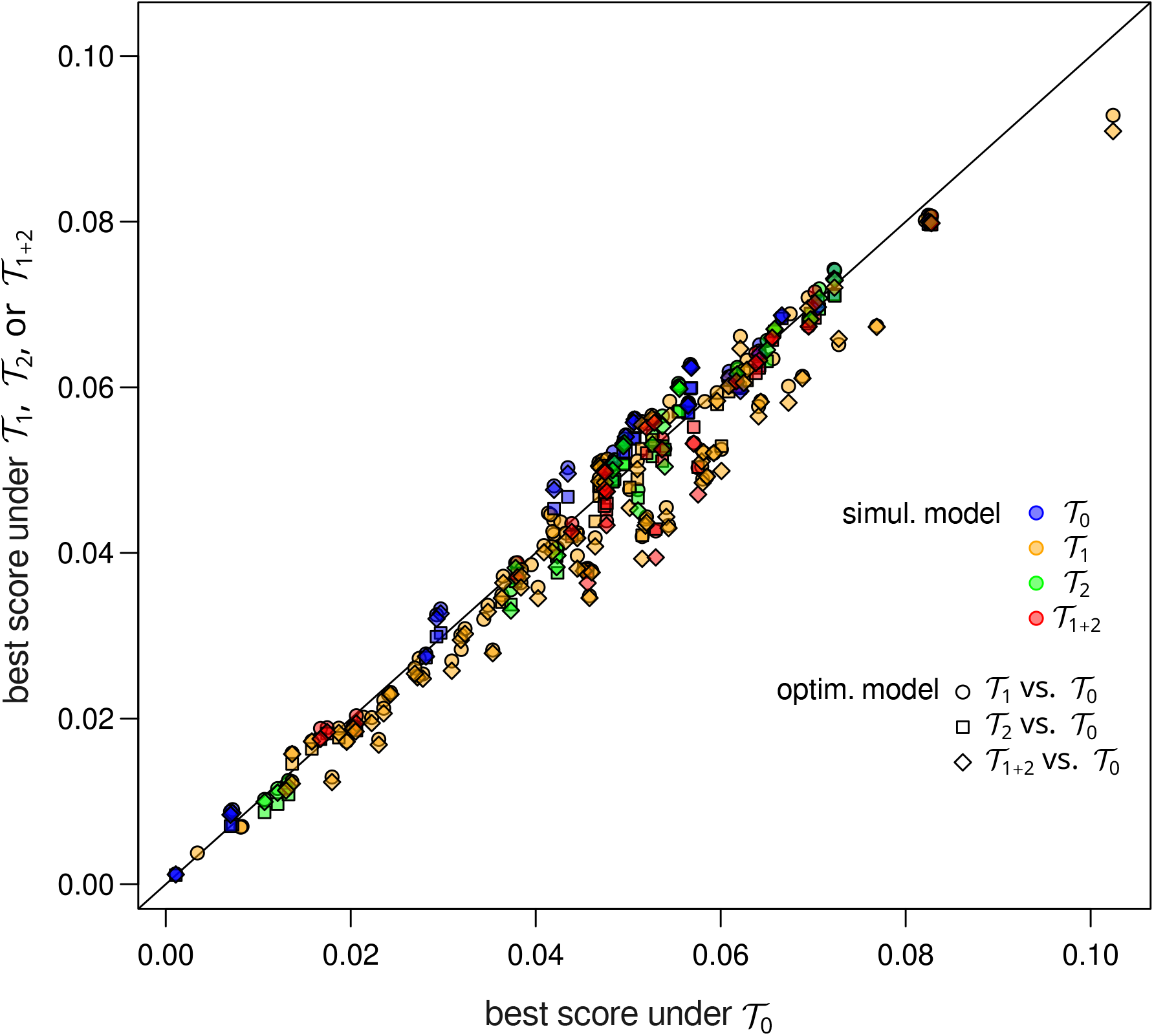
Comparison of the scores obtained with different transcription models on the same simulated datasets. Comparison between the best score (score *p*^*⋆*^ of the best-scoring path) obtained under optimization with one of the three models for transcription initiation ( T_1_, T_2_, or T_1+2_) and the best score obtained under model T_0_ which assumes a negligible effect of transcription initiation on the occupancy ratios. The model considered for the impact of transcription initiation is indicated by the shape of the symbol; the model used for the simulation of the dataset is indicated by the color of the symbol. The points corresponding to dataset simulated with T_0_ tends to be located above the first diagonal, whereas the points corresponding to datasets simulated with T_1_, T_2_, and T_1+2_, tends to be located below the first diagonal.

**Figure S12:**
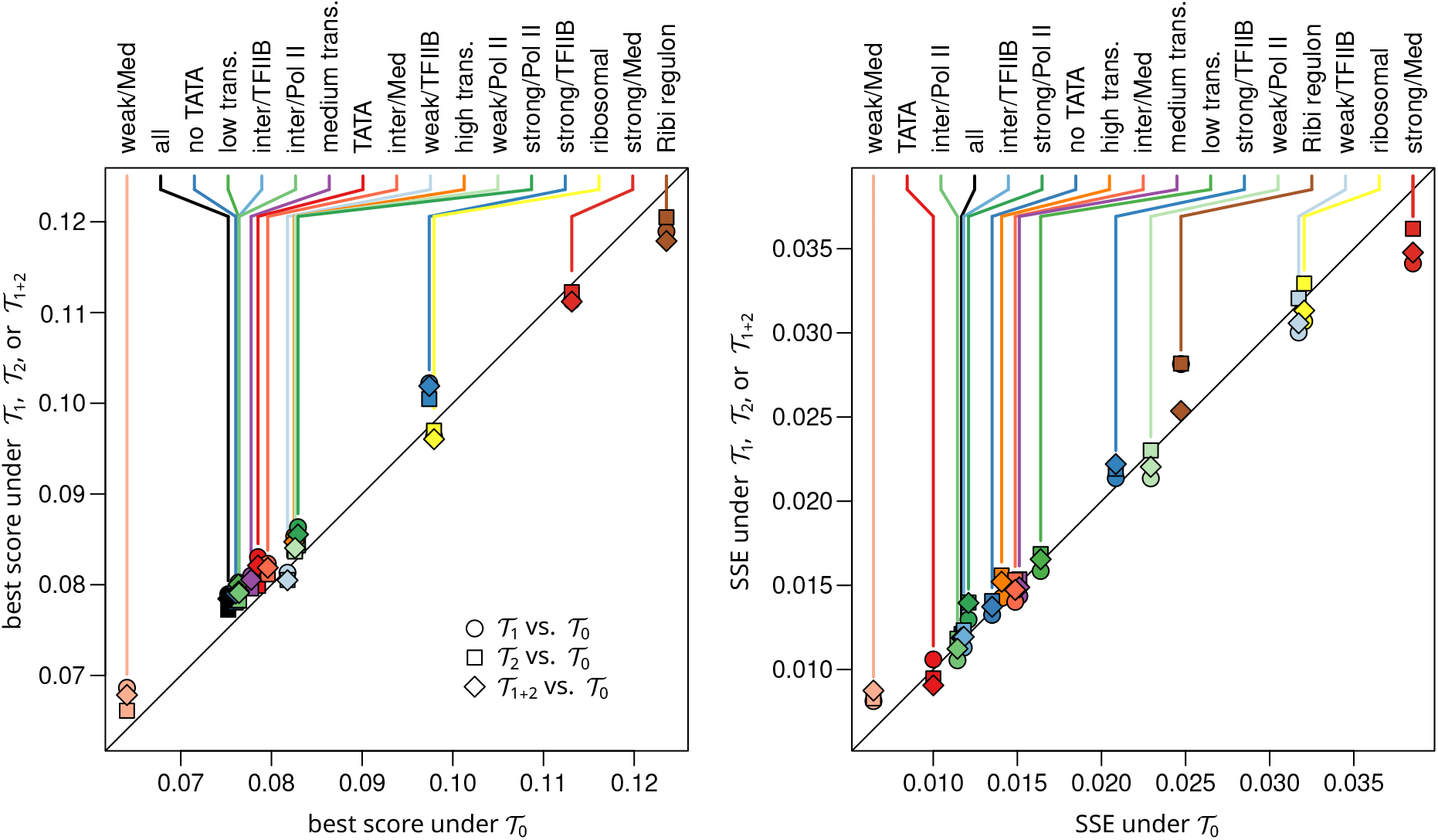
Comparison between transcription models across the different groups of genes. Each point corresponds to the analysis of the mutant vs. wild-type profile for one of the 17 considered groups of genes. Left panel: comparison between the best score (score *p*^*⋆*^ of the best-scoring path) under model T_1_, T_2_, or T_1+2_, (y-axis) versus the best score under model T_0_ (x-axis). The symbols distinguish the model with transcription whose score is represented on the y-axis: circles for model T_1_, squares for model T_2_, lozenges for model T_1+2_. Right panel: comparison between the sum of squared errors of prediction for the best-scoring path. Penalization coefficients: *λ* = 10^*−*1.5^ and *µ* = 10^*−*2^.

**Figure S13:**
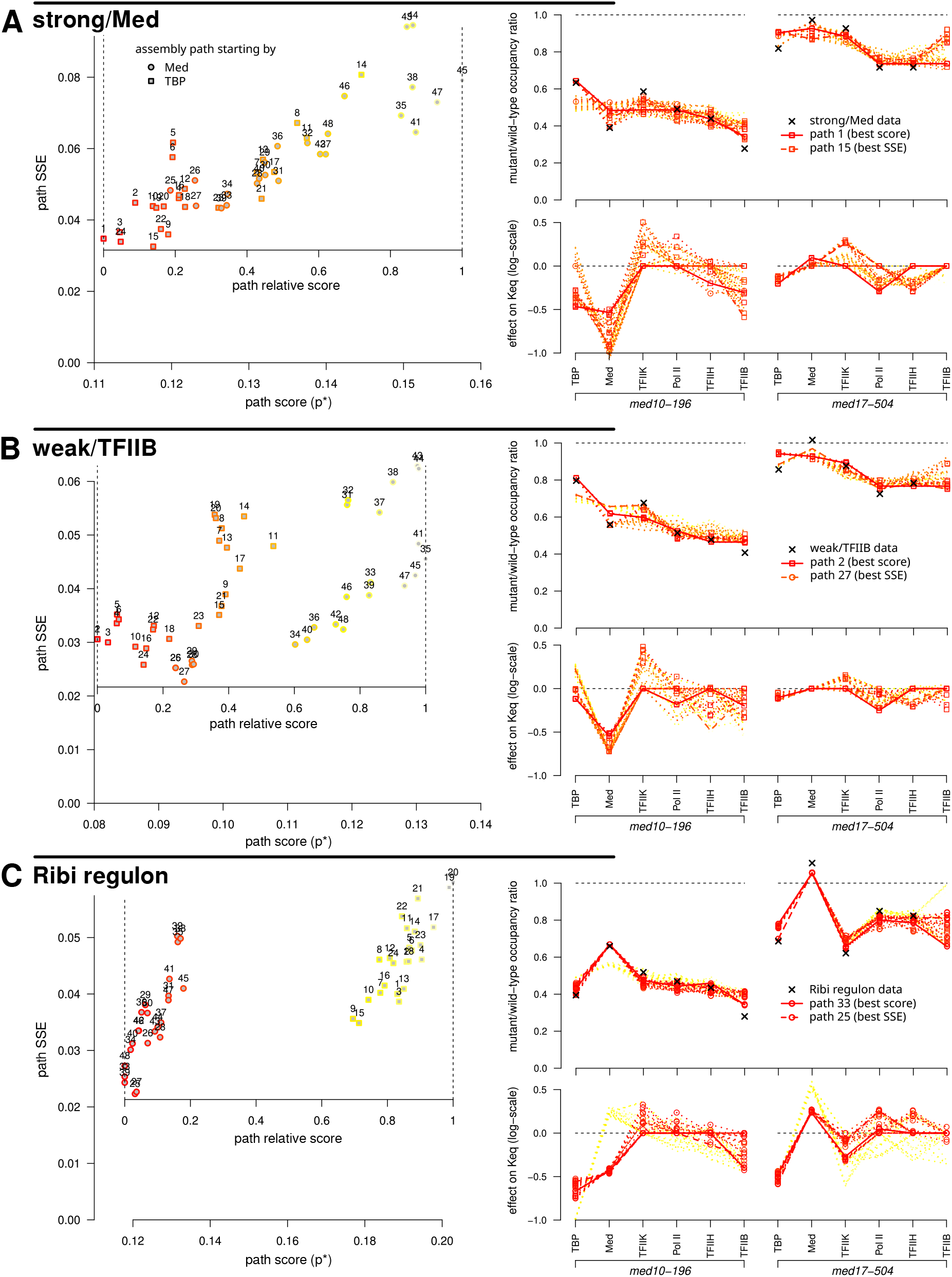
Path scores, goodness of fit and estimated parameters for the profiles of three subgroups of genes. **A** Results for the mutant vs. wild-type occupancy profile of the group “strong/Med”, *i*.*e*. the most impacted genes with respect to Med occupancy. **B** Results for the mutant vs. wild-type occupancy profile of the group “weak/TFIIB”, *i*.*e*. the least impacted genes with respect to TFIIB occupancy. **C** Results for the mutant vs. wild-type occupancy profile of the group “Ribi regulon”. See Figure 6 for the legend of the subpanels within panels **A-C**. The results presented here were obtained with model T_1+2_, since the best scores are achieved with this model for the profiles of these three groups (Supplementary Fig. S12), instead of T_0_, as used in Figure 6. Penalization coefficients: *λ* = 10^*−*1.5^ and *µ* = 10^*−*2^.

